# Short prokaryotic Argonautes provide defence against incoming mobile genetic elements through NAD^+^ depletion

**DOI:** 10.1101/2021.12.14.472599

**Authors:** Mindaugas Zaremba, Donata Dakineviciene, Edvardas Golovinas, Evelina Zagorskaitė, Edvinas Stankunas, Anna Lopatina, Rotem Sorek, Elena Manakova, Audrone Ruksenaite, Arunas Silanskas, Simonas Asmontas, Algirdas Grybauskas, Ugne Tylenyte, Edvinas Jurgelaitis, Rokas Grigaitis, Kęstutis Timinskas, Česlovas Venclovas, Virginijus Siksnys

**Affiliations:** Institute of Biotechnology, Life Sciences Center, Vilnius University, Saulėtekio av. 7, LT-10257, Vilnius, Lithuania; Department of Molecular Genetics, Weizmann Institute of Science, Rehovot 7610001, Israel

## Abstract

Argonaute (Ago) proteins are found in all three domains of life. The so-called long Agos are composed of four major domains (N, PAZ, MID, and PIWI) and contribute to RNA silencing in eukaryotes (eAgos) or defence against invading mobile genetic elements in prokaryotes (pAgos). The majority (~60%) of pAgos identified bioinformatically are shorter (comprised of only MID and PIWI domains) and are typically associated with Sir2, Mrr or TIR domain-containing proteins. The cellular function and mechanism of short pAgos remain enigmatic. Here, we show that *Geobacter sulfurreducens* short pAgo and the NAD^+^-bound Sir2-protein form a stable heterodimeric complex. The GsSir2/Ago complex presumably recognizes invading plasmid or phage DNA and activates the Sir2 subunit, which triggers endogenous NAD^+^ depletion and cell death, and prevents the propagation of invading DNA. We reconstituted NAD^+^ depletion activity *in vitro* and showed that activated GsSir2/Ago complex functions as a NADase that hydrolyses NAD^+^ to ADPR. Thus, short Sir2-associated pAgos provide defence against phages and plasmids and underscores the diversity of mechanisms of prokaryotic Agos.

---

Being at the core of RNA interference eAgos are involved in the regulation of gene expression, silencing of mobile genome elements, and defence against viruses^1,2^. The best-studied hAgo2 uses small RNA molecules as guides for target RNA recognition, and eAgos are similar both structurally and mechanistically^2–5^. Monomeric eAgos are composed of four major N, PAZ (<u>P</u>iwi/<u>A</u>rgonaute/<u>Z</u>wille), MID (<u>Mid</u>dle), and PIWI (<u>P</u>-element <u>I</u>nduced <u>Wi</u>mpy testis) domains (Fig. 1A) and share a bilobed structure, where the N- and C-terminal lobes are formed by conserved N/PAZ and MID/PIWI domains, respectively^4–7^. The N-domain acts as a wedge that separates guide and target strands^3,7^, while the MID and PAZ domains bind, respectively, the 5’- and 3’-terminus of the guide RNA, located between the N- and C-terminal lobes^4,6^. eAgos can slice target RNA through endonucleolytic cleavage by the PIWI domain or inhibit translation through RNA binding by the catalytically inactive eAgos that may also trigger RNA decay by auxiliary cellular nucleases^2,4,5^.

**Fig. 1.**
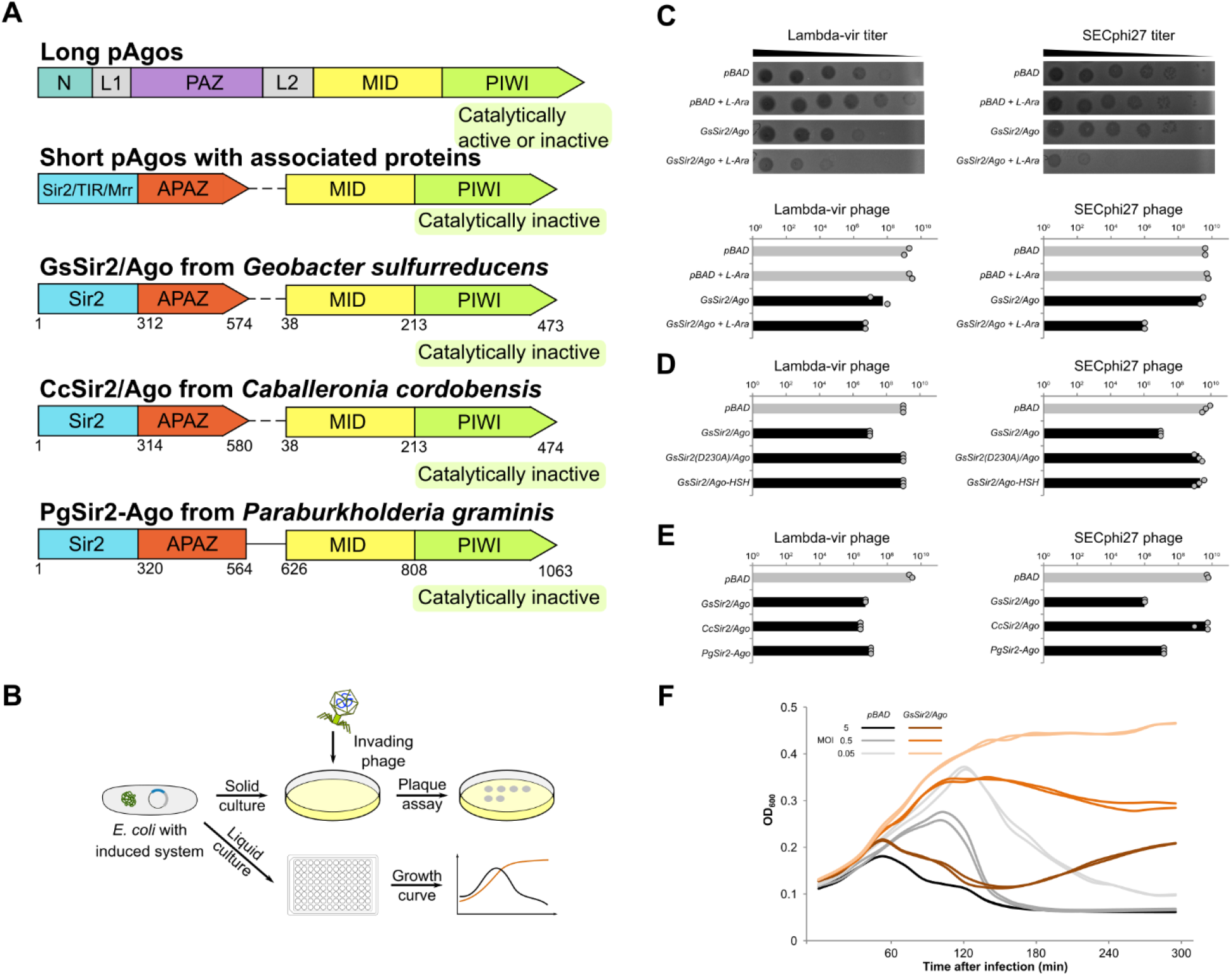
Sir2/Ago provide defence against phages. **A**, Schematic representation of the subunit/domain composition of different pAgo variants. Catalytically active pAgos contain a conserved catalytic DEDX tetrad that is mutated in inactive pAgos. Catalytically inactive short pAgos lack the canonical PAZ domain, however, an accessory APAZ domain is present in putative Sir2, TIR or Mrr proteins associated with short pAgos. MID indicates middle; L – linker domain; N – N-terminal domain. Short pAgos from *G. sulfurreducens, C. cordobensis*, and *P. graminis* associated with Sir2 protein were studied in this work. **B**, Schematic diagram of phage restriction assays. **C**, Efficiency of plating (EOP) of lambda-vir and SECphi27 phages infecting *E. coli* cells with and without the GsSir2/Ago system. The bar graphs show the number of p.f.u. as arithmetic means of two replicates in the absence and the presence of the inducer L-arabinose (L-Ara), with individual data points superimposed. Grey bars represent EOP on pAgo-lacking cells and black bars – in pAgo-containing cells. Representative images of plaque assays are also presented. **D**, EOP of lambda-vir and SECphi27 phages infecting the wt and mutant GsSir2/Ago systems in the presence of L-Ara. GsSir2(D230A)/Ago and GsSir2/Ago-HSH are variants that contain D230A mutation in the Sir2 domain or (HSH-tag) on the C-terminus of pAgo, respectively. The bar graphs show the number of p.f.u. as arithmetic means of three replicates, with individual data points superimposed. Grey bars represent EOP on pAgo-lacking cells and black bars – in pAgo-containing cells. **E**, EOP of lambda-vir and SECphi27 phages infecting pAgo-lacking cells and cells containing GsSir2/Ago, CcSir2/Ago and PgSir2-Ago in the presence of L-Ara. The bar graphs show the number of p.f.u. as arithmetic means of three replicates, with individual data points superimposed. Grey bars represent EOP on pAgo-lacking cells and black bars – in pAgo-containing cells. **F**, Lambda phage infection in liquid cultures of *E. coli* cells containing the GsSir2/Ago system. GsSir2/Ago-lacking (shown in grey) or GsSir2/Ago-containing (shown in orange) *E. coli* were infected at t=0 at multiplicities of infection (MOI) of 0.05, 0.5 and 5. Each curve represents one individual replicate; two replicates for each MOI are shown.

pAgos are quite widespread and are present in 9% of sequenced bacterial and 32% of archaeal genomes^6^. To date, more than ~1000 pAgos have been identified bioinformatically, revealing a striking diversity. Moreover, pAgos are often associated with additional putative nucleases, helicases, and DNA binding proteins that are not linked to eAgos^8^. pAgos are divided into full-length or long pAgos (~40%) sharing conserved N, PAZ, MID, and PIWI domain architecture with eAgos, and short pAgos (~60%) composed only of MID and PIWI domains (Fig. 1A)^8,9^. Long pAgos are relatively well-characterized both structurally and functionally^10,11^ and, similarly to eAgos, contain either catalytically active or inactive PIWI domain. Some long pAgos with the catalytically active PIWI, as exemplified by CbAgo and TtAgo, use DNA guides to target and cleave DNA providing defence against invading phages or plasmids, or contributing to chromosome segregation after replication, respectively^12–14^. Meanwhile, long RsAgo with an inactive PIWI domain guided by small RNA is thought to mobilize an unknown cellular nuclease(s) for degradation of invading plasmids and mobile genetic elements^9,15^. Interestingly, long KmAgo can use both DNA and RNA guides to target DNA and RNA *in vitro*, albeit with different efficiencies^16,17^. In contrast to long pAgos, all short pAgos possess a catalytically inactive PIWI domain and are typically associated with proteins containing a domain initially thought to be analogous to PAZ (APAZ)^18^. Subsequently, however, it was proposed that APAZ may actually be homologous to the N-terminal domains (N and L1) of Ago^11,19^. APAZ-containing proteins are often fused to Sir2 (<u>S</u>ilent <u>i</u>nformator <u>r</u>egulator <u>2</u>), Mrr nucleases or TIR (<u>T</u>oll-<u>I</u>nterleukin-1 <u>R</u>eceptor) domains^8,18^. About half of all identified short pAgos are associated or fused into a single-chain protein with Sir2-APAZ proteins^8^. The Sir2 domain-containing proteins are widely distributed in all domains of life and perform protein deacetylation or ADP-ribosylation functions using NAD^+^ as a co-factor^20,21^. Particularly, bacterial Sir2 proteins are involved in many cellular processes including transcription, translation, carbon and nitrogen metabolism, virulence and resistance to stress^21^. Despite the fact that short pAgos, half of which are associated with Sir2 proteins, make up the majority of all pAgos, their function in the cell and *in vitro* remains to be established.

## Results

In this study, we aimed to explore whether short pAgos can act as prokaryotic defence systems against viruses or plasmids. To this end, we selected two short pAgos, GsSir2/Ago from *Geobacter sulfurreducens* and CcSir2/Ago from *Caballeronia cordobensis*, each encoded in a putative operon together with a Sir2 domain protein, and PgSir2-Ago from *Paraburkholderia graminis*, representing a fusion of Sir2 and pAgo (Fig. 1A). The coding regions of the Sir2 and pAgo proteins in GsSir2/Ago and CcSir2/Ago systems overlap by 11 and 8 bp, respectively, indicating that they belong to the same operon (Supplementary Text). Next, we engineered heterologous *E. coli* cells by cloning GsSir2/Ago and CcSir2/Ago genes, or a single gene, encoding PgSir2-Ago into pBAD expression vectors under a P_BAD_ promoter (Supplementary Table 1) and challenged them with phages or plasmids.

### Sir2/Ago systems provide defence against phages

To test whether the GsSir2/Ago system provides defence against phages, we challenged *E. coli* host carrying the GsSir2/Ago system with a set of six *E. coli* phages spanning four morphological families including *Podoviridae* (T7), *Siphoviridae* (lambda-vir, SECphi27, SECphi18), *Myoviridae* (T4) and *Microviridae* (SECphi17, a ssDNA phage) (Supplementary Table 1). We measured the efficiency of plating (EOP) of these phages with and without L-arabinose induction of the GsSir2/Ago system. The system showed protection against two out of six phages – lambda-vir (~100-fold) and SECphi27 (~1000-fold) (Fig. 1C).

To probe the role of individual Sir2 and Ago proteins in antiviral defence, we engineered *E. coli* cells, carrying mutant variants of either the Sir2 or the Ago protein of the GsSir2/Ago system and performed small drop plaque assays using the lambda-vir and SECphi27 phages. In Sir2 variants, the highly-conserved D230 residue, presumably involved in NAD^+^ binding was replaced by alanine (Extended Data Fig. 1C)^18^. Phage challenge assay revealed that the D230A mutation completely abolished defence against both phages (Fig. 1D). The Ago protein is catalytically inactive due to the active site mutations in the PIWI domain, therefore, to obtain a binding-deficient Ago variant, we fused a bulky 29 aa **H**is_6_-**S**trepII-**H**is_6_-tag (HSH-tag) at the C-terminus that is important for nucleic acid binding in other Agos^3,6,8,22^. We found that Ago C-terminal modification abolished protection from both phages (Fig. 1D). Thus, both the Sir2 and the Ago proteins are required for protection against phages by the GsSir2/Ago system.

Additionally, we probed the ability of homologous CcSir2/Ago and PgSir2-Ago systems to restrict lambda-vir and SECphi27 phages (Fig. 1E). After L-arabinose induction, the PgSir2-Ago system showed ~500-fold protection against lambda phage and ~400-fold against SECphi27, while CcSir2/Ago system showed ~1000-fold protection against lambda and no protection against SECphi27 phage. Thus, homologous Sir2/Ago systems from three phylogenetically distant bacteria showed protection against phage infection, albeit with different efficiency and specificity.

Next, *E. coli* MG1655 cells either lacking or containing the GsSir2/Ago system were infected in liquid culture with lambda phage at a multiplicity of infection (MOI) 0.05, 0.5 and 5 (Fig. 1F). At high MOI where, on average, a single bacterial cell is infected by a single phage, the culture collapses, while at low MOI the culture survives. This phenotype implies that GsSir2/Ago-mediated defence triggers cell death approximately at the same time when the phage-induced lysis occurs.

### Sir2/Ago systems interfere with plasmid transformation

Further, to test whether heterologous expression of the GsSir2/Ago, CcSir2/Ago and PgSir2-Ago systems in *E. coli* cells (strains BL21-AI and DH10B) provides a barrier for plasmid transformation, four plasmids (pCDF, pCOLA, pACYC184, and pRSF) with different *ori* regions and copy numbers were used in a plasmid interference assay (Supplementary Table 1, Fig. 2, and Extended Data Fig. 2). We found that GsSir2/Ago system prevented only pCDF plasmid transformation reducing its efficiency nearly ~100-fold (Fig. 2B). Next, we tested pCDF transformation efficiency in *E. coli* cells expressing the GsSir2/Ago mutants (Fig. 2C). Both Sir2 D230A mutation and pAgo HSH-tag modification that impaired phage restriction also abolished plasmid interference making cells permissive for pCDF plasmid transformation (Fig. 2C). CcSir2/Ago system like GsSir2/Ago provided resistance only for pCDF plasmid transformation (Extended Data Fig. 2B) while cells carrying the single-chain PgSir2-Ago system was permissive for transformation of all four plasmids (Extended Data Fig. 2B).

**Fig. 2.**
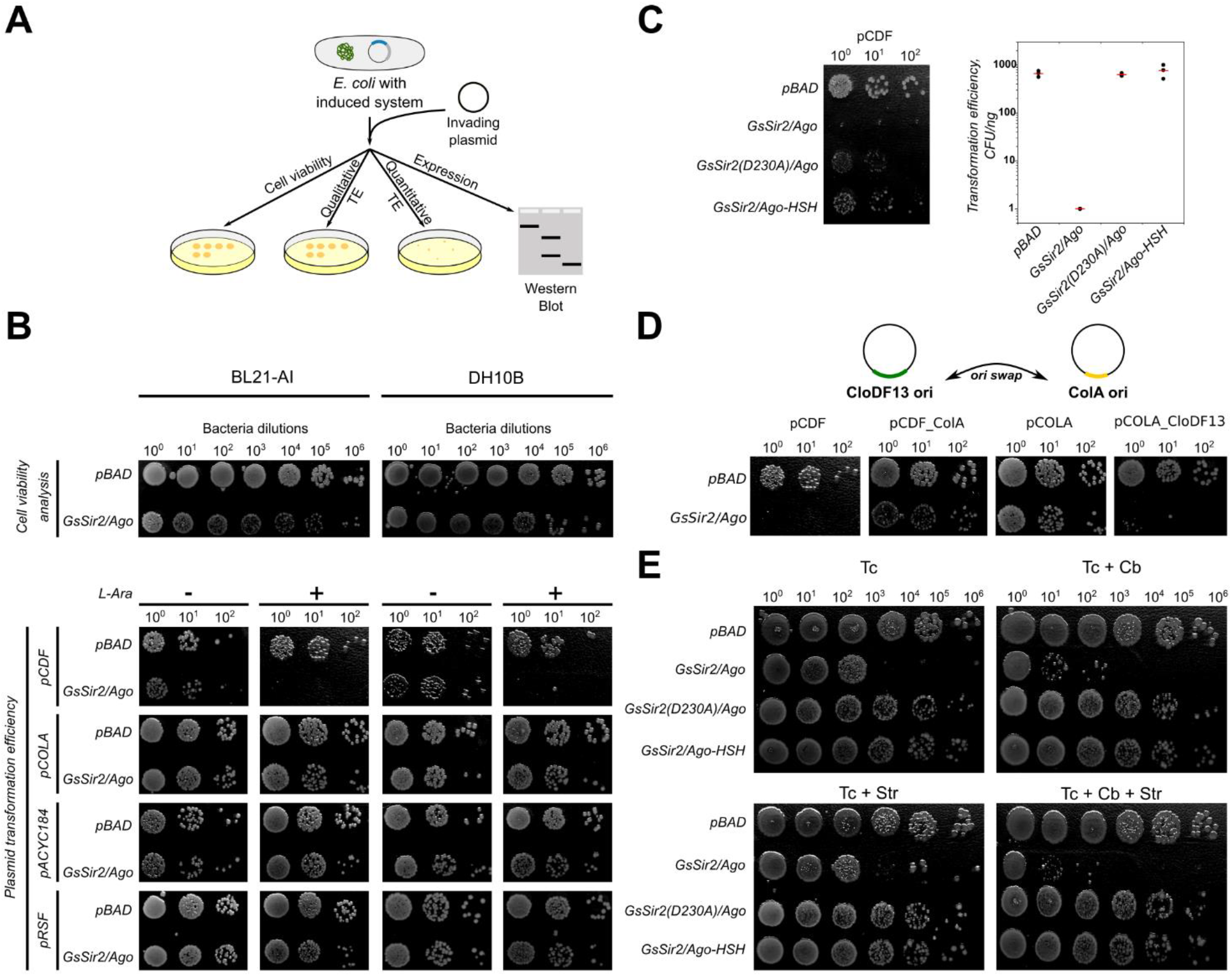
The GsSir2/Ago system interferes with plasmid transformation. **A**, Schematic representation of the experiment. **B**, Qualitative evaluation of plasmid transformation efficiency in *E. coli* cells carrying GsSir2/Ago system. Top: comparison of cell viability in the presence or absence of plasmid-borne GsSir2/Ago expression. Bottom: comparison of plasmid transformation efficiencies in the presence or absence of plasmid-borne GsSir2/Ago expression. **C**, Left – comparison of pCDF transformation efficiency between cells expressing wt and mutant GsSir2/Ago complexes; right - quantification of transformation efficiencies (three independent replicates, the red line represents average transformation efficiency). **D**, Top - schematic representation of *ori* exchange between pCDF and pCOLA plasmids; bottom - comparison of plasmid transformation efficiencies. pCDF, pCDF with CloDF13 *ori* exchanged with ColA *ori* (pCDF_ColA), pCOLA and pCOLA with ColA *ori* exchanged with CloDF13 *ori* (pCOLA_CloDF13) plasmids were used for transformation of *E. coli* cells carrying GsSir2/Ago system. **E**, Cell viability in the absence of antibiotic selection. In the case of the wt GsSir2/Ago system, the cell viability decreases on the plates even in the absence of Cb and Str antibiotics suggesting that GsSir2/Ago in the presence of the pCDF plasmid triggers cell death.

Interestingly, although the GsSir2/Ago system in *E. coli* interfered with the pCDF plasmid transformation, pCOLA plasmid that differs mainly in the *ori* region and antibiotic resistance gene was permissive. To test whether the *ori* region determines differences in the transformation efficiency between pCDF and pCOLA plasmids, we swapped the *ori* sequences of pCDF and pCOLA (Fig. 2D). The pCDF plasmid with ColA *ori* instead of CloDF13 became permissive in *E. coli* cells expressing the GsSir2/Ago system, whereas transformation of pCOLA plasmid bearing CloDF13 *ori* instead of ColA *ori* was prevented. These results indicate that CloDF13 *ori* is a key element that controls plasmid transformation efficiency in *E. coli* cells expressing GsSir2/Ago system.

The plasmid interference by the GsSir2/Ago system could be due to either the plasmid entry exclusion, replication inhibition or plasmid degradation. To eliminate the possible role of the plasmid entry barriers on the pCDF plasmid transformation efficiency, we engineered heterologous *E. coli* cells carrying two plasmids: pBAD plasmid providing carbenicillin (Cb) resistance and expressing GsSir2/Ago (or its mutants) under control of P_BAD_ inducible promoter and the pCDF plasmid providing streptomycin (Str) resistance, and tested cell viability in the presence or absence of the inducer. In this case, the pCDF plasmid is already in the cell and provides streptomycin resistance, however antibiotic resistance should be lost if the plasmid is restricted after GsSir2/Ago expression. In the absence of induction, cell viability of *E. coli* cells carrying wt GsSir2/Ago (or its mutants) and an empty pBAD vector (Extended Data Fig. 2E) was identical. In the presence of the inducer the viability of cells expressing the wt GsSir2/Ago system, but not its mutants, significantly decreased (Fig. 2E), indicating that GsSir2/Ago interferes with the pCDF plasmid already present in the cell. Notably, a decrease in cell viability is observed in *E. coli* BL21-AI cells (Tc-resistant) without the cell selection for Str and Cb resistance. It cannot be excluded that upon recognition of pCDF the GsSir2/Ago system becomes activated and triggers cell death. A similar cell death phenotype triggered by the GsSir2/Ago has been observed during phage infection in liquid cultures (Fig. 1F). Taken together, these data show that the GsSir2/Ago system acts as a defence system against phages and plasmids via cell death or suicidal mechanism.

### Short pAgo and Sir2 form a stable heterodimeric complex

To characterize the Sir2/Ago systems biochemically, we aimed to express individual Sir2 and Ago proteins in *E. coli*. The GsSir2 and CcSir2 proteins (but not PgSir2-Ago) were expressed and purified by chromatography (Fig. 3A and Extended Data Fig. 3). The N-terminal His_6_-tagged GsSir2 and CcSir2 proteins co-expressed with Ago proteins co-purified on the Ni^2+^-affinity column (Extended Data Fig. 3), indicating that Sir2 and pAgo proteins form a stable complex. We failed to express Sir2 and Ago proteins individually, suggesting that they form an obligatory Sir2/Ago complex. Sir2/Ago complex. Functionally compromised GsSir2(D230A)/Ago and GsSir2/Ago-HSH variants also formed a complex indicating that while the introduced mutations abolished the activity *in vivo*, they did not affect the protein complex structure (Fig. 2C, Extended Data Fig. 3C). Further analysis of the oligomeric state of GsSir2/Ago and CcSir2/Ago complexes in solution using SEC-(MALS), mass photometry and small-angle X-ray scattering (SAXS) showed that heterodimeric complexes are formed in the wide range of protein concentrations (from 20 nM to 6.5 μM) (Fig. 3B-D, and Extended Data Fig. 4). According to the SAXS data, the heterodimeric GsSir2/Ago complex acquires a notably asymmetric shape (Fig. 3D, Extended Data Fig. 4, Supplementary Table 3) that is consistent with a structural model of the heterodimer (Supplementary Text). In summary, the results show that pAgos and associated Sir2 proteins encoded by a single operon form a stable heterodimeric complex.

**Fig. 3.**
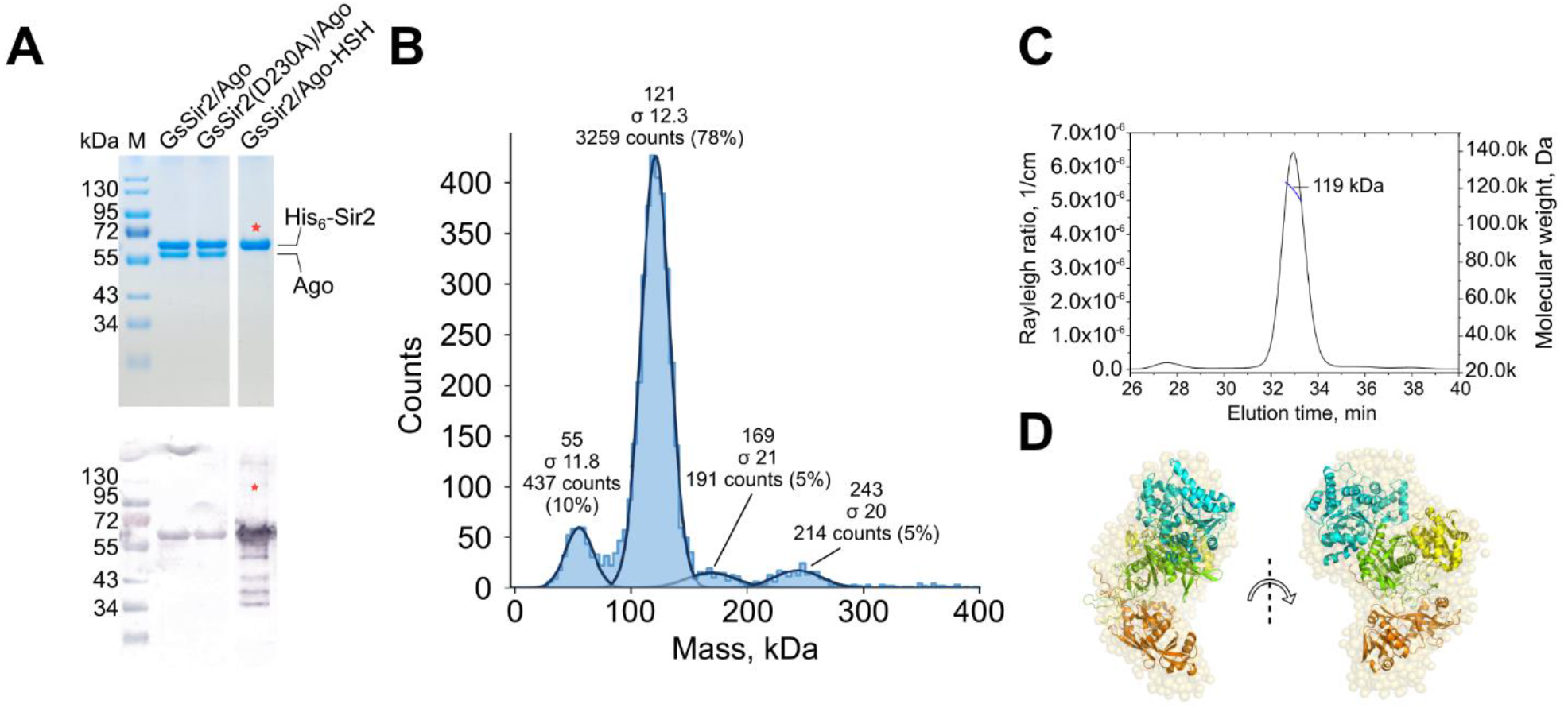
The GsSir2 and GsAgo proteins form a heterodimeric complex. **A**, Top – SDS PAGE of purified wt GsSir2/Ago, a D230A mutant, and C-terminal HSH tag-bearing GsSir2/Ago. Red star in the HSH-tagged sample lane marks an overlap of bands in the gel due to similar mass. Bottom – anti-His-tag Western blot of same samples. The red star shows the lane where the His-tag is on the C-terminus of Ago, rather than the N-terminus of Sir2. Two replicates. **B**, Mass photometry data of the GsSir2/Ago complex, with masses and respective particle population counts indicated. According to mass spectrometry of the purified GsSir2/Ago complex (Extended Data Fig. 3F), the molar mass of the GsSir2/Ago heterodimer is 121 kDa. **C**, SEC-MALS data of GsSir2/Ago, showing the chromatography peak and molar weight of the Sir2/Ago heterodimer. **D**, Semitransparent space-filling *ab initio* model of GsSir2/Ago calculated from SAXS data with a fitted-in AF GsSir2/Ago model in cartoon representation. Colour coding: Sir2 domain – cyan, APAZ – brown, MID – yellow, PIWI – green.

### Sir2/Ago prefers an RNA guide to bind a DNA target

Long pAgos use ssDNA and/or ssRNA guides to recognize their complementary DNA and/or RNA targets^10^. To establish guide preference of Gs and Cc Sir2/Ago, we analysed by EMSA binding of single- or double-stranded DNA or RNA (Extended Data Fig. 6). Both Sir2/Ago heterodimers showed a strong preference for ssDNA and ssRNA binding. RNA/DNA heteroduplex was bound with an intermediate affinity, while dsRNA or dsDNA showed only weak binding (Extended Data Fig. 6A-B,E, Table 1). Interestingly, neither the 5’-terminal phosphate nor the 3’-OH end or Mg^2+^ ions were required for ssDNA binding as both Gs and Cc complexes bound the circular ssDNA and linear oligonucleotides with similar affinity (Extended Data Fig. 6A-B). As expected, Gs and Cc Sir2/Ago containing the inactivated PIWI domains showed no cleavage activity for any NA substrate tested (Extended Data Fig. 6F). In summary, EMSA experiments suggest that *in vitro* ssRNA or ssDNA are preferable GsSir2/Ago guides.

**Table 1.** Nucleic acid binding by Sir2/Ago and their binary complexes pre-loaded with either ssRNA or ssDNA guides. K_d_ values (averages ± standard deviation of three independent replicates) were calculated from EMSA data. Some binding experiments were performed in the presence of heparin (indicated).

| Nucleic acid binding by Sir2/Ago |  |  |  |
| --- | --- | --- | --- |
| Nucleic acid | | $K_d$, nM | |
|  |  | GsSir2/Ago | CcSir2/Ago |
| ssDNA | | $0.033 \pm 0.016$ | $0.138 \pm 0.007$ |
| ssRNA | | $0.042 \pm 0.015$ | $0.36 \pm 0.098$ |
| circular ssDNA | | $0.072 \pm 0.018$ | $0.18 \pm 0.068$ |
| dsRNA/DNA | | $4.31 \pm 0.37$ | $2.4 \pm 0.69$ |
| dsDNA | | $51 \pm 20$ | $330 \pm 87$ |
| dsRNA | | $73 \pm 29$ | $81 \pm 15$ |
| Nucleic acid binding by GsSir2/Ago-gRNA and GsSir2/Ago-gDNA complexes |  |  |  |
| Guide | Target | $K_d$, nM | |
|  |  | - Heparin | + Heparin |
| ssRNA | ssDNA | $0.020 \pm 0.008$ | $0.079 \pm 0.007$ |
| | ssRNA | $0.205 \pm 0.058$ | $0.567 \pm 0.112$ |
| ssDNA | ssDNA | $0.022 \pm 0.005$ | n.b.* |
| | ssRNA | $0.252 \pm 0.114$ | n.b.* |
| ssRNA | nsp-ssDNA<sup>#</sup> | $0.420 \pm 0.231$ | $4.29 \pm 2.11$ |
| ssDNA binding by GsSir2/Ago-gRNA variants |  |  |  |
| Mutant | | $K_d$, nM | |
|  |  | - Heparin | + Heparin |
| GsSir2(D230A)/Ago | | $0.038 \pm 0.013$ | $0.140 \pm 0.057$ |
| GsSir2/Ago-HSH | | $0.140 \pm 0.017$ | $0.203 \pm 0.058$ |
| GsSir2<sup>APAZ</sup>/Ago |  | n.b.* | n.b.* |
| GsSir2/Ago<sup>PIWI</sup> |  | n.b.* | n.b.* |

\* No binding was detected under experimental conditions used. <sup>#</sup> A non-complementary ssDNA was used.

Next, we analysed ssDNA or ssRNA target binding by the binary GsSir2/Ago complexes pre-loaded with either ssRNA or ssDNA guides. In a separate set of experiments, reaction mixture also contained heparin, a competitor of nucleic acid binding (Fig. 4A, Table 1). The binary GsSir2/Ago-ssRNA complex showed ~10-fold better binding to the ssDNA than ssRNA, and heparin addition had only a little effect (~4-fold decrease) on binding affinity in this case. The GsSir2/Ago-ssDNA binary complex bound to the matching ssDNA target with affinity similar to the GsSir2/Ago-ssRNA binary complex, however, heparin addition abolished binding (Extended Data Fig. 6A and D). Furthermore, the binary GsSir2/Ago-ssRNA complex bound to the complementary ssDNA target ~200-fold better than apo-GsSir2/Ago bound pre-annealed gRNA/tDNA heteroduplex indicating that GsSir2/Ago requires binding of the RNA guide first to interact with the DNA target (Fig. 4A, Table 1). The binding affinity of the functionally compromised *in vivo* GsSir2(D230A)/Ago mutant was similar to the wt, while the binding affinity of the GsSir2/Ago-HSH mutant was slightly (~7-fold) weaker (Fig. 4A, Extended Data Fig. 6C, Table 1). Charge reversal mutations of positively charged residues that are involved in the interactions with the guide and target NAs according to the GsSir2/Ago model abolished both the target NA binding and the pCDF plasmid interference (Extended Data Fig. 2F and G, Extended Data Fig. 3B and Extended Data Fig. 6C). Taken together, our data suggest that GsSir2/Ago uses ssRNA as a guide for the recognition of a ssDNA target.

**Fig. 4.**
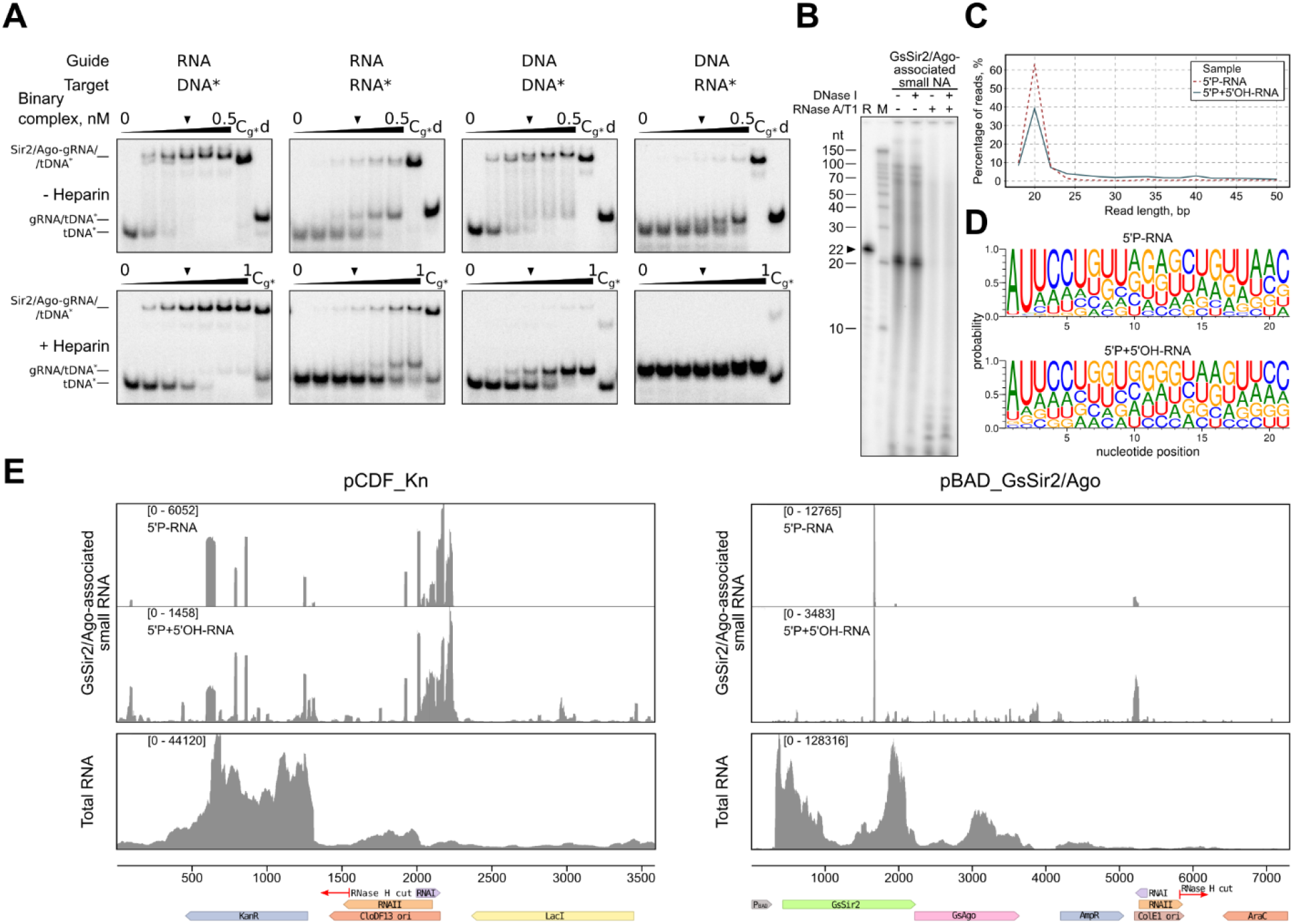
Nucleic acid binding by GsSir2/Ago *in vitro* and *in vivo*. **A**, Binding of RNA or DNA targets by GsSir2/Ago binary complexes pre-loaded with 5’P-RNA or 5’P-DNA guides. In EMSA experiments the pre-formed GsSir2/Ago-gNA binary complex was mixed with a radiolabelled target strand indicated by the asterisk (see “Materials and Methods” for the details). To show that no displacement of the guide by the target strand occurs under these experimental conditions, a control (Cg*) experiment was performed where the guide, rather than the target, was radioactively labelled. Only the pre-annealed RNA/DNA heteroduplex was loaded in the control lane “d”. Three independent replicates performed. **B,** GsSir2/Ago co-purifies with small RNAs. Nucleic acids that co-purified with GsSir2/Ago were, first, dephosphorylated, then, [γ-^32^P]-ATP radiolabelled and treated with DNase I or RNase A/T1, or both, and resolved on a denaturing polyacrylamide gel. R, control 22 nt RNA oligonucleotide; M, RNA ladder Decade Marker System (Ambion). Three independent replicates performed. **C**, Length distribution of small RNA co-purified with GsSir2/Ago as determined by sequencing. “5’P-RNA” sample means that only small RNAs containing 5’-phosphate were sequenced, while in “5’P+5’OH-RNA” sample both 5’-phosphate or 5’-OH bearing small RNAs were sequenced. **D**, Small RNAs associated with GsSir2/Ago show 5’-AU preference. **E**, Distribution of small RNAs co-purified with GsSir2/Ago from the *E. coli* host carrying the pCDF_Kn target and pBAD_GsSir2/Ago expression plasmids (above). IGV viewer representation of total RNA extracted from *E. coli* is shown below. Cartoons indicate promoters (P_BAD_), protein-coding genes (KanR, LacI, GsSir2, GsAgo, AmpR, AraC), plasmid *ori* (CloDF13, ColE1) and their RNAI and RNAII transcripts. Red arrow shows RNase H cleavage site in RNAII required for initiation of DNA strand synthesis during plasmid replication.

To identify NAs, bound by GsSir2/Ago *in vivo*, we purified the GsSir2/Ago-NA complex from *E. coli* transformed with the pBAD_GsSir2/Ago expression vector and the pCDF target plasmid, extracted NAs and subjected them to sequencing. Subsequent analysis revealed that GsSir2/Ago is associated with small (predominantly 21 nt) RNAs with or without the 5’-phosphate (Fig. 4B and C). Differently from other Argonaute proteins that show base selectivity for the first nucleotide at the 5’-end of the guide^13–15,23^, GsSir2/Ago-associated small RNAs show preference for the 5’-AU dinucleotide (Fig. 4D). This preference is more pronounced for small RNAs containing 5’-phosphate, implying that GsSir2/Ago uses as a guide small RNAs containing the 5’-AU dinucleotide. Most co-purified small RNAs (~95%) matched the *E. coli* genome, whereas the smaller fraction (~5%) originated from the pBAD_GsSir2/Ago and pCDF plasmids (Supplementary File 1). Interestingly, small plasmid-borne RNAs that matched CloDF13 and ColE1 *ori* regions of corresponding plasmids, were noticeably enriched (Fig. 4E). Taken together, RNA-seq data suggest that GsSir2/Ago could use small 5’-AU-RNAs originating from the invader transcripts (e.g., pCDF *ori* region) as guides to target the invaders’ DNA.

### The GsSir2/Ago complex binds NAD^+^ and causes its depletion

Computational analysis of Sir2 domains showed that they possess a conserved NAD^+^-binding pocket (Extended Data Fig. 1C and Extended Data Fig. 5). To determine whether Sir2 domains can indeed bind endogenous NAD^+^, purified GsSir2/Ago and CcSir2/Ago complexes were heat-treated, protein aggregates removed by centrifugation, and the supernatant analysed by MS-HPLC (Fig. 5 and Extended Data Fig. 7). The quantitative analysis showed that both the GsSir2/Ago and CcSir2/Ago complexes co-purified with bound endogenous NAD^+^ in approx. 1:1 molar (Sir2:NAD^+^) ratio (Fig. 5A, Extended Data Fig. 3, Extended Data Fig. 7). However, in the case of the functionally inactive GsSir2(D230A)/Ago mutant, only 0.6% of all complexes were NAD^+^-bound indicating that the mutation severely compromised NAD^+^ binding by the Sir2 domain (Fig. 5A). NAD^+^ binding by the GsSir2 subunit and the similarity of the GsSir2 to the N-terminal NADase of the ThsA protein from the anti-phage Thoeris system^24^ prompted us to investigate the level of endogenous NAD^+^ in the presence of the induced GsSir2/Ago system and its target pCDF plasmid. In these experiments the corresponding *E. coli* cells were lysed, proteins were removed, and the amount of NAD^+^ in the supernatant was examined by MS-HPLC. When the wt GsSir2/Ago expression was induced in the presence of pCDF plasmid, NAD^+^ was depleted (~120-fold decrease), whereas in the case of the functionally inactive GsSir2(D230A)/Ago and GsSir2/Ago-HSH mutants the level of endogenous NAD^+^ was similar to that of the empty pBAD vector (Fig. 5B, Extended Data Fig. 7). It should be noted that a significant ~30-fold decrease of the NAD^+^ level was observed when the wt GsSir2/Ago expression was induced even in the absence of pCDF suggesting that the heterologously expressed system may be toxic to the cells resulting in their slower growth (Fig. 5B, Extended Data Fig. 7). To test the hypothesis that the activated GsSir2/Ago system similarly to the Thoeris anti-phage system depletes the endogenous NAD^+^ through hydrolysis or cyclization, we attempted to identify possible products (ADPR, cADPR, AMP, cAMP, ADP, cADP, nicotinamide, and adenine) using MS-HPLC albeit without success. It is possible that NAD^+^ conversion products were not detected since in the cell they were processed to other reaction intermediates or due to ion suppression during MS analysis of the cell lysates.

**Fig. 5.**
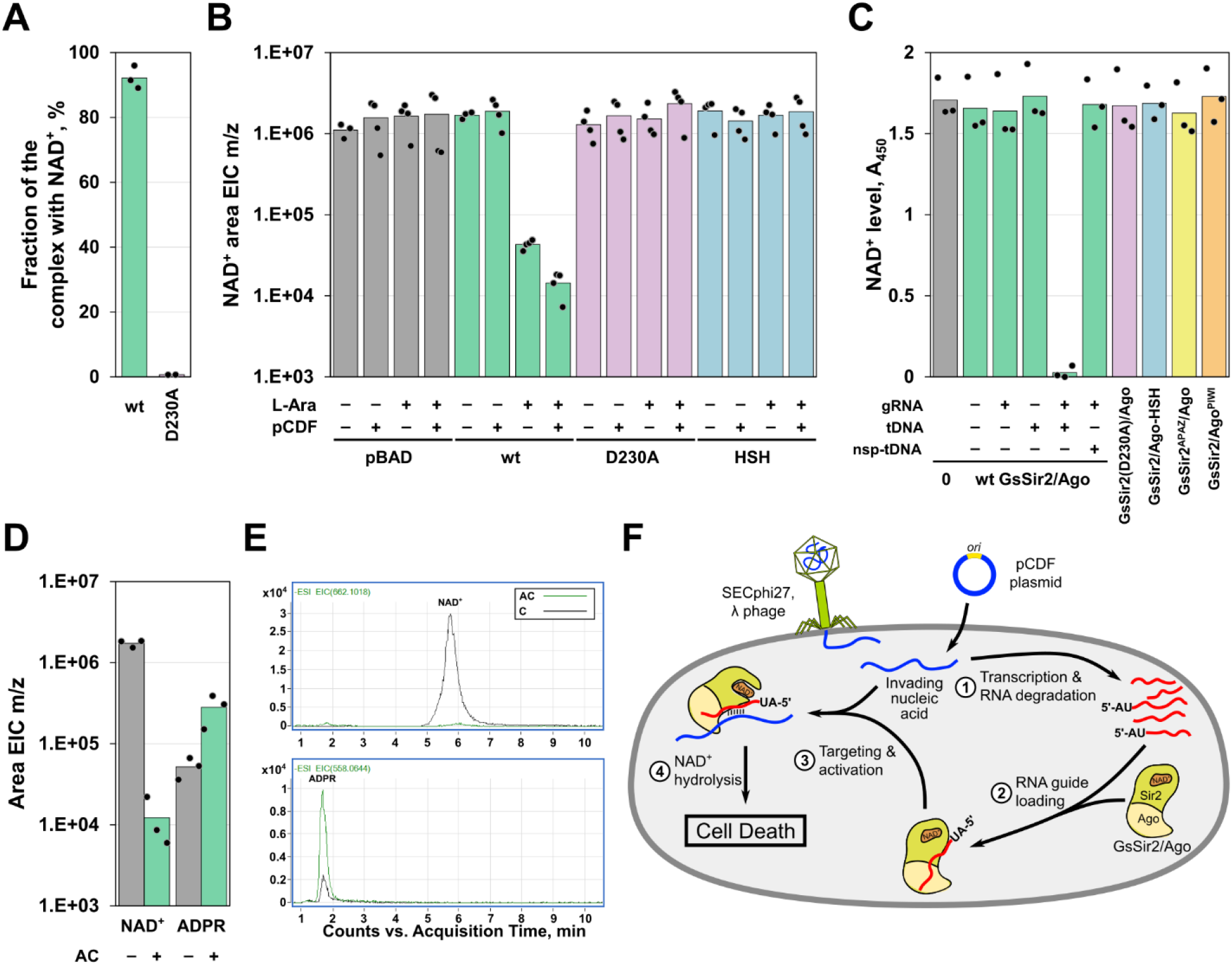
GsSir2/Ago binds and hydrolyses NAD^+^. **A**, the D230A mutation within the Sir2 protein abolished NAD^+^ binding. **B**, NAD^+^ amounts in *E. coli* cells in the presence of the (non)induced wt and mutant GsSir2/Ago systems and in the presence and absence of the pCDF plasmids. NAD^+^ amounts were estimated according to the EIC areas of NAD^+^ (m/z 662.1018). pBAD – empty vector; wt – GsSir2/Ago; D230A – GsSir2(D230A)/Ago; HSH – GsSir2/Ago-HSH. **C**, NAD^+^ depletion by GsSir2/Ago *in vitro*. The wt GsSir2/Ago or mutant complex (0.5 μM) was incubated with NAD^+^ (50 μM) for 1 h at 37 °C (see experimental details in “Materials and Methods”). gRNA, 5’P-RNA guide; tDNA, target DNA complementary to the RNA guide; nsp-tDNA, ssDNA non-complementary to the RNA guide. **D**, NAD^+^ hydrolysis by wt GsSir2/Ago *in vitro*. The binary GsSir2/Ago-gRNA complex (0.5 μM) was incubated with NAD^+^ (50 μM) for 1 h at 37 °C in the presence of the complementary DNA target (0.5 μM). NAD^+^ depletion and ADPR accumulation was analysed by MS according to the EIS of NAD^+^ (m/z 662.1018) and ADPR (m/z 558.0644), respectively. AC, the activated wt GsSir2/Ago-gRNA/tDNA complex. **E**, Representative mass chromatograms of NAD^+^ hydrolysis by wt GsSir2/Ago *in vitro* (as in **D**). C, a control sample without GsSir2/Ago; AC, the activated wt GsSir2/Ago-gRNA/tDNA complex. **F**, Putative model of GsSir2/Ago defence against mobile genetic elements. After lambda phage infection or pCDF plasmid transformation, GsSir2/Ago acquires small 5’-AU-RNAs originated from the invader transcripts (e.g., from pCDF *ori* region). The GsSir2/Ago binary complex, guided by small RNA, targets the invaders’ complementary DNA, becomes activated and hydrolyses NAD^+^ resulting in a cell death.

Next, we investigated whether the NAD^+^ depletion activity detected in cells could be reconstituted *in vitro*. To this aim we mixed the binary GsSir2/Ago-gRNA complex with ssDNA and monitored NAD^+^ concentration using a commercial kit (Fig. 5C). We found that NAD^+^ concentration decreased when wt GsSir2/Ago-gRNA complex was added to the complementary ssDNA target, however, no changes were observed in the case of non-complementary ssDNA. Mutations in the Sir2 domain (D230A) or the APAZ/Ago part (GsSir2/Ago-HSH, GsSir2^APAZ^/Ago and GsSir2/Ago^PIWI^ mutants) compromised NAD^+^ depletion (Fig. 5C). MS analysis revealed that GsSir2/Ago-gRNA complex in the presence of the complementary ssDNA hydrolyses NAD^+^ to ADPR (Fig. 5D,E) similarly to the Thoeris anti-phage system^24^. Taken together, these results demonstrate that GsSir2/Ago functions as a NADase that becomes activated upon target DNA binding.

## Discussion

Association of Sir2-like domains with short pAgo proteins has been identified bioinformatically in the pioneering Makarova et al. paper^18^. It has been speculated that Sir2-domain proteins can act as nucleases, however, the structure and function of Sir2 proteins so far have not been elucidated. Here, we show that the APAZ-containing Sir2 and short pAgo proteins form a heterodimeric complex (Fig. 3) similarly as in the case of a short pAgo and a Mrr nuclease domain-containing protein^25^. Furthermore, our structure modelling results show that the APAZ region of Sir2 proteins shares similarity with the N, L1 and L2 domains of canonical Agos substantiating previous sequence-based predictions^11,19^ At the same time, Sir2 proteins entirely lack the PAZ domain (Fig. 1A, Extended Data Fig. 5A). Thus, apparently both split and single-chain Sir2/Ago systems evolved from long pAgos by the loss of the PAZ domain and acquisition of the Sir2 domain.

Next, we provide the experimental evidence that the Sir2/Ago complex functions as a defence system against invading phages and plasmids (Fig. 1 and Fig. 2). Intriguingly, plasmid interference assay using four plasmids with different replicons (Extended Data Fig. 2) revealed that GsSir2/Ago system prevents transformation only of the pCDF plasmid that contains CloDF13 *ori* (Fig. 2), suggesting that GsSir2/Ago may recognize specific replicon elements or structures. Indeed, *ori* swap between pCDF and permissive pCOLA plasmid made the latter sensitive to GsSir2/Ago interference. We show here that GsSir2/Ago co-purifies from *E. coli* cells together with small (predominantly 21 nt long) RNAs that preferentially contain the 5’-AU dinucleotide (Fig. 4B-D). Interestingly, a fraction of small RNAs that originates from pCDF CloDF13 and pBAD ColE1 *ori* regions is enriched (Fig. 4E). ColE1-like origins, including CloDF13, use two small RNAs (RNAI and RNAII) for priming of the replication that involves the R-loop intermediate^26–28^. It is tempting to speculate that GsSir2/Ago is able to bind nucleic acids of different length (Fig. 4B,C) and preferentially binds *ori-*associated small RNAs that can be subjected to further processing by cellular RNases to produce ~21 nt gRNAs similarly to long RsAgo that shares an inactivated PIWI domain with GsSir2/Ago^15^.

We further show that *in vitro* the reconstituted wt GsSir2/Ago-gRNA complex becomes activated after binding the complementary DNA target and triggers NAD^+^ hydrolysis generating ADPR (Fig. 5C-E). It is likely, that in the *E. coli* cells the APAZ/Ago part in the GsSir2/Ago complex guided by the *ori-*associated RNA guides could bind to the complementary plasmid or phage DNA target activating the Sir2 effector domain that depletes endogenous NAD^+^ leading to the cell death, thereby restricting plasmid and phage propagation (Fig. 5F, Supplementary text, the accompanying paper by Garb et al., 2021^29^). A similar anti-phage defence mechanism based on NAD exhaustion has been shown for the Thoeris and the Pycsar systems, CBASS (<u>C</u>yclic Oligonucleotide-<u>Ba</u>sed <u>S</u>ignaling <u>S</u>ystem) and DSR (<u>D</u>efense-<u>A</u>ssociated <u>S</u>irtuins)^29–31^. In the Thoeris system of *Bacillus cereus* MSX-D12, the Sir2 domain is similar to that of the GsSir2/Ago system and performs the hydrolysis of NAD^+^ to ADPR and nicotinamide^31^ like the GsSir2/Ago system. Further structural and biochemical studies are underway to establish the structure of the heterodimeric Sir2/Ago complex and the mechanism of Sir2 domain activation that triggers NAD^+^ hydrolysis.

## Methods

### Oligonucleotides used in this work

All synthetic DNA oligonucleotides used for cloning and site-specific mutagenesis were purchased from Metabion (Germany) and are listed in Supplementary Table 2.

### Cloning and mutagenesis

A whole operon of the GsSir2/Ago system, composed of the Sir2 (GSU1360, NP_952413.1) and Ago (GSU1361, NP_952414.1) encoding genes, was amplified by PCR from the genomic DNA of *Geobacter sulfurreducens* Caccavo (ATCC No. 51573, LGC Standards cat#51573D-5) using the oligonucleotides MZ-239 and MZ-240 (Supplementary Table 2), respectively. The resulting DNA fragment was digested by Eco31I (ThermoFisher cat#FD0293) and XhoI (ThermoFisher cat#FD0694) and cloned into pBAD/HisA expression vector (ThermoFisher cat#V43001) pre-cleaved with NheI (ThermoFisher cat#FD0973) and XhoI and dephosphorylated using FastAP (ThermoFisher cat#EF0651). The D230A mutant of the GsSir2 protein was produced by QuikChange Site-Directed Mutagenesis^32^ using respective mutagenic oligonucleotides (Supplementary Table 2). To generate the GsAgo protein containing a bulky **H**is_6_-**S**trepII-**H**is6-tag (HSH-tag, 29 aa.: LEGHHHHHHSSWSHPQFEKGVEGHHHHHH) a whole operon of the GsSir2/Ago system was amplified by PCR from the genomic DNA using the oligonucleotides MZ-325 and MZ-326 (Supplementary Table 2), respectively. The resulting DNA fragment was digested by Eco31I and XhoI cloned into pBAD24 vector through NcoI/XhoI sites to generate pBAD24-HSH expression vector. The GsSir2/Ago mutants (GsSir2^APAZ^/Ago, GsSir2/Ago^MID^ and GsSir2/Ago^PIWI^) of the putative surface of the interaction with nucleic acids were designed based on the GsSir2/Ago structural model (see, Supplementary methods) changing positively charged residues that are structurally equivalent to the RsAgo residues involved in the interaction with the guide and the target to negatively charged residues in the corresponding domains (APAZ: R440E/R442E/R517E/R525E/R528E/R531E, MID: K155E/H166E/K170E/K207E, PIWI: R269E/K270E/H310E/K441E). The surface mutants were obtained using whole gene synthesis and cloning service provided by Twist Bioscience.

A PgSir2-Ago gene (BgramDRAFT_6510, WP_052303232.1) was amplified by PCR from the genomic DNA of *Paraburkholderia graminis* C4D1M (ATCC No. 700544) purchased from LGC Standards (UK) using the oligonucleotides MZ-915 and MZ-916 (Supplementary Table 2), respectively. The resulting DNA fragment was digested by BveI (ThermoFisher cat#FD1744) and HindIII (ThermoFisher cat#FD0504) and ligated into pBAD24 expression vector precleaved with NcoI and HindIII and dephosphorylated using FastAP.

The *E. coli* codon optimized genes (IDT Codon Optimization Tool) encoding the Sir2 (WP_053571900.1) and Ago (WP_053571899.1) of the CcSir2/Ago system (from *Caballeronia cordobensis*, NCBI taxon_id 1353886) were synthesized and cloned into pBAD/HisA expression vector by Twist Bioscience. The CcSir2 protein contains at its N-terminus a His_6_-tag that can be cleaved by TEV protease. For purification of the CcSir2/Ago complex, a TwinStrep-tag (35 aa.: MGGSAWSHPQFEKGGGSGGGSGGSAWSHPQFEKGS) was additionally fused to the N-terminus of the CcSir2 protein already containing a His_6_-tag.

To swap *ori* regions between pCOLA and pCDF plasmids the DNA fragments containing ColA and CloDF13 *ori* were amplified by PCR using the oligonucleotides MZ-1217/MZ-1218 and MZ-1230/MZ-123, respectively (Supplementary Table 2). The resulting DNA fragments were digested by NheI and XbaI (ThermoFisher cat#FD0684) and ligated into pCOLA and pCDF vectors precleaved with NheI and XbaI and dephosphorylated using FastAP.

To swap streptomycin resistance for kanamycin in the pCDF plasmid, plasmids pCDF and pCOLA were cleaved with NheI and Eco81I (ThermoFisher cat#FD0374) and isolated using a runView electrophoresis system (Cleaver Scientific). The purified fragments were then ligated into pCDF to yield a pCDF_Kn plasmid.

All gene sequences were confirmed by sequencing; links to DNA and protein sequences are presented in Supplementary Table 1.

### Phage restriction assay

*E. coli* MG1655 (ATCC 47076) cells carrying pBAD plasmids expressing wt or mutated Sir2/Ago systems were used for phage infection assays as described below. Whole-plasmid sequencing was applied to all transformed *E. coli* clones to verify the integrity of the system and lack of mutations, as described before^33^.

*E. coli* phages (T4, T7, lambda-vir) were kindly provided by U. Qimron. Phages SECphi17, SECphi18 and SECphi27 were isolated by the Sorek lab as described before^34^. Small drop plaque assay was performed as described by Mazzocco et al.^35^. Overnight culture of *E. coli* bacteria was diluted 1:100 in MMB medium (LB + 0.1 mM MnCl_2_ + 5 mM MgCl_2_ + 5 mM CaCl_2_) supplied with 0.1% L-arabinose for expression induction. Bacterial cultures were incubated at 37 °C until early log phase (OD_600_ = 0.3), and 500 μl of bacteria were mixed with 25 ml of MMB agar (LB + 0.1 mM MnCl_2_ + 5 mM MgCl_2_ + 5 mM CaCl_2_ + 0.5% agar + 0.1% L-arabinose) and poured into the square Petri dish. Serial dilutions of phage lysate in MMB were dropped on top of cell lawn. After the drops dried up, plates were incubated at room temperature for 24 hours. Efficiency of plating (EOP) was determined via comparison of phage lysate titer on control bacteria and bacteria containing the Argonaute system with and without induction with L-arabinose.

Liquid culture phage infection experiments were performed as described previously by Ofir et al.^33^. After overnight incubation, the liquid suspension of pAgo-lacking and pAgo-containing *E. coli* cells were diluted 1:100 in MMB medium supplied with 0.2% L-arabinose and dispensed into a 96-well plate by 180-μl volume. Plates were incubated at 37 °C until the early log phase (OD_600_ = 0.3), then 20 μl of phage lysate was added to each well in multiplicity of infection 5, 0.5, or 0.05, and each experiment was performed in three replicates. Optical density measurements at a wavelength 600 nm were taken every 15 min using a TECAN Infinite 200 plate reader.

### Plasmid interference assay

*E. coli* BL21-AI and DH10B strain cells were pre-transformed with pBAD/HisA plasmid encoding a GsSir2/Ago (either wild-type or mutant variants), CcSir2/Ago or PgSir2-Ago - under the control of araBAD promoter. After 2 hours of induction with either 0.01% (w/v) (CcSir2/Ago) or 0.1% (GsSir2/Ago and PgSir2-Ago) L-arabinose at 37 °C 200 RPM, cells were heat-shock transformed with pCDF (pCDF_Kn in the case of PgSir2-Ago in DH10B), pCOLA, pACYC184 or pRSF plasmids. After recovery, cells were either serially diluted and aliquots spotted on selection medium or undiluted suspensions spread on selection medium and CFUs counted manually. In parallel, viability and over-expression of pAgo-containing *E. coli* cells were monitored using serial dilutions and Western blot, respectively.

In separate experiments, *E. coli* BL21-AI strain cells were heat-shock transformed with both GsSir2/Ago operon-encoding pBAD/HisA construct and pCDF plasmid. Protein expression in selected double transformants was then induced by the addition of L-Ara (final concentration of 0.1% (w/v)) into liquid LB culture. After a 2-hour induction at 37 °C 200 RPM, optical density (OD_600_) equalized, cultures were serially diluted and aliquots were spotted on a selection medium containing different antibiotics.

### Expression and purification of GsSir2/Ago complexes

For GsSir2/Ago protein expression *E. coli* DH10B strain was transformed with a respective plasmid (Supplementary Table 1). Cells were grown at 37 °C in LB medium in the presence of 50 μg/ml ampicillin until OD_600_ = 0.7 was reached. Then, the temperature was decreased to 16 °C and proteins were expressed for 16 h by adding 0.2% w/v L-arabinose. Next, harvested cells were disrupted by sonication in buffer A (20 mM Tris–HCl (pH 8.0 at 25 °C), 500 mM NaCl, 2 mM phenylmethylsulfonyl fluoride, 5 mM 2-mercaptoethanol), and cell debris was removed by centrifugation. GsSir2/Ago complexes were purified to >90% homogeneity by chromatography through HisTrap HP chelating, HiTrap Heparin HP and HiLoad Superdex 200 columns (GE Healthcare). Purified proteins were stored at −20 °C in a buffer containing 20 mM Tris-HCl (pH 8.0 at 25 °C), 200 mM KCl, 1 mM DTT and 50% v/v glycerol. The identity of the purified proteins was confirmed by mass spectrometry. Protein concentrations were determined from OD_280_ measurements using the theoretical extinction coefficients calculated with the ProtParam tool available at http://web.expasy.org/protparam/. GsSir2/Ago complex concentrations are expressed in terms of heterodimer. The GsSir2/Ago^MID^ surface mutant could not be purified due to its poor expression.

### Antibodies used in this work

For Western blot analysis of target proteins, the following antibodies were used: 6x-His Tag monoclonal antibody (ThermoFisher, cat. #MA1-21315, RRID AB_557403, Lot # WE323793, clone HIS.H8); Goat anti-Mouse IgG (H+L) Secondary Antibody, AP conjugated (ThermoFisher, cat. #31320, RRID AB_228304, Lot # VH311913, polyclonal).

### SEC-(MALS) and mass photometry

Size-exclusion chromatography of GsSir2/Ago complexes was carried out at room temperature using Superdex 200 10/300 GL column (GE Healthcare) pre-equilibrated with a buffer (20 mM Tris–HCl (pH 8.0 at 25 °C), 500 mM NaCl). A calibration curve was generated by measuring the elution volumes of a series of standard proteins of known molecular mass (Bio-Rad). The molecular masses of pAgos complexes were calculated by interpolating their elution volume onto the calibration curve. SEC-MALS of GsSir2/Ago and CcSir2/Ago complexes was performed at room temperature using Superdex 200 10/300 GL column (GE Healthcare) pre-equilibrated with a buffer (20 mM Tris–HCl (pH 8.0 at 25 °C), 500 mM NaCl, 0.03% NaN3, 1 mM DTT), at 0.4 ml/min flow rate. Sample concentrations were 6 μM and 6.5 μM for GsSir2/Ago and CcSir2/Ago, respectively. The light scattering signals were monitored on a miniDawn TREOS II detector, concentrations of protein samples were measured using an Optilab T-rEX refractive index detector (Wyatt Technologies). Data were analysed in Astra software (Wyatt Technologies) using dn/dc value of 0.185 mL/g. Mass photometry of the GsSir2/Ago complex was performed using a Refeyn OneMP system (Refeyn). The protein complex was diluted to 20 nM in a buffer containing 20 mM Tris-HCl pH 8.0, 500 mM NaCl before measurement.

### Small-angle X-ray scattering (SAXS) analysis

The synchrotron SAXS data were collected at beamline P12 operated by EMBL Hamburg at the PETRA III storage ring (DESY, Hamburg, Germany)^36^. GsSir2/Ago sample in the storage buffer was transferred into the sample buffer (20 mM Tris-HCl pH7.5, 200 mM NaCl, 5 mM MgCl_2_, 2 mM β-mercaptoethanol) using gel-filtration NAP column (GE Healthcare) and concentrated by ultrafiltration to 1.2, 1.3, 1.6 and 5.5 mg/ml concentrations. The data were collected at the wavelength of 0.124 nm and the distance to the detector (Pilatus 2M, Dectris) was set to 3 m. Samples in the sample changer were kept at 10 °C, capillary temperature was set to 20 °C. Twenty frames exposed 0.045 sec were averaged for each concentration. The s-range of collected data was from 0.0133796 to 3.7925 nm^-1^. The data were analysed using programs of ATSAS 2.8.4 (r10552) suite^37^. Data were normalized to an absolute scale with water as standard. As the data collected for the sample with a concentration of 1.22 mg/ml were noisy at higher s (Extended Data Fig. 4), and higher concentration data showed more aggregation at low s, we used a merged dataset produced with PRIMUS^38^. Scattering data were parameterized and indirectly Fourier transformed with GNOM5^39^. Structural parameters of this dataset are summarized in Supplementary Table 3. Dimensionless Kratky plot in Extended Data Fig. 4 was calculated as described previously^40^. The *ab initio* models were calculated by GASBOR^41^ software. Molecular mass estimations of apo GsSir2/Ago complex in solution assessed by ATSAS tools (DATVC, DATMW) and server SAXSMoW (http://saxs.ifsc.usp.br/)^42^ are presented in the Supplementary Table 3.

### Nucleic acid binding assay

The oligonucleotide substrates (Supplementary Table 2) were 5’-labelled with [γ-^32^P]ATP (PerkinElmer) and T4 polynucleotidyl kinase (PNK) (ThermoFisher cat#EK0031). The 3’-labelled substrate was prepared with [α-^32^P]cordycepin-5’-triphosphate (Hartmann Analytics) and terminal deoxynucleotidyl transferase. An aliquot of the 3’-labelled substrate was subsequently phosphorylated with cold ATP and T4 PNK to obtain a 3’[α-^32^P]cordycepin-labelled oligonucleotide containing 5’-phosphate. Annealing was performed in the PNK reaction buffer supplemented with 50 mM EDTA at 2 μM total single-stranded oligonucleotide concentration. Circular ssDNA substrate was prepared by circularisation of 5’-^32^P labelled TK-49 using CircLigase II (Lucigen cat#CL9021K), according to manufacturer recommendations, and purified from a denaturing PAA gel (21% 29:1 acrylamide/bis-acrylamide in TBE supplemented with 8 M urea) by phenol-chloroform extraction, precipitated in 96 % ethanol with 0.45 M sodium acetate, washed with 75% ethanol, and resuspended in water.

For EMSA experiments, appropriate substrates and proteins were pre-diluted to 2x the final binding reaction concentration in 40 mM Tris-acetate (pH 8.3 at 23 °C), 1 mM EDTA (TAE, Invitrogen cat#24710-030), supplemented with 5 mM magnesium acetate, 0.1 mg/ml BSA, 1 mM DTT, and 10% glycerol. The binding reactions were conducted by mixing equal volumes of enzyme and radiolabelled substrate. In all cases, final binding reactions contained 0.1 nM of radiolabelled substrate at 0-2 nM (0; 0.02; 0.05; 0.1; 0.2; 0.5; 1; 2) or 0-500 nM (0; 5; 10; 20; 50; 100; 200; 500) of GsSir2/Ago and CcSir2/Ago complexes. Three independent replicates were performed. For clarity, only EMSA gels obtained using low protein concentrations (0-2 nM) were shown, while for calculations of high K_d_ values EMSA data obtained using high protein concentrations (0-500 nM) were used.

Binding experiments for the binary GsSir2/Ago complex were conducted by first pre-mixing 5’-phosphorylated ssRNA or ssDNA guide with the equimolar GsSir2/Ago complex in the same buffer as above. The binary GsSir2/Ago:NA guide complex was then diluted to 2x final reaction concentration (in respect to guide) in the same buffer and mixed with a complementary 5’-^32^P-target oligonucleotide in the presence or absence of 67 ng/μl heparin sodium salt (SigmaAldrich cat#). The final reaction contained 10 pM target NA and 0, 0.02, 0.05, 0.1, 0.2 0.5, and 1 nM of GsSir2/Ago:NA guide complex. Control (Cg*) contained 0.1 nM GsSir2/Ago:NA guide complex with the guide labelled with [γ-^32^P]ATP and 10 pM unlabeled target NA. Three independent replicates were performed.

The binding reaction mixtures were analysed by electrophoretic mobility shift assay (EMSA) in a PAA gel (8% 29:1 acrylamide/bis-acrylamide in TAE). The electrophoresis TAE buffer was supplemented with 5 mM magnesium acetate. Radiolabelled substrates were detected and quantified using a phosphor imager. The results were analysed with OptiQuant and OriginPro software. The K_d_ was calculated from the following formula:

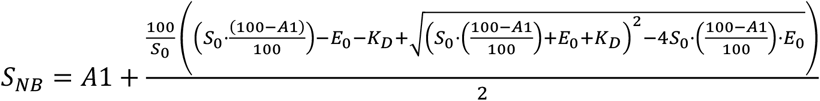

where S_NB_ – unbound substrate, nM; S_0_ – initial substrate concentration, nM; E_0_ – initial protein complex concentration, nM; K_d_ – dissociation constant, A1 – nonbinding fraction of substrate, %.

### Nucleic acid extraction and analysis

To obtain GsSir2/Ago-bound nucleic acids, *E. coli* DH10B was transformed with pBAD/HisA_TwinStrep_TEV_GsSir2/Ago and pCDF_Kn plasmids (Supplementary Table 1). Cells were grown at 37 °C in LB medium in the presence of 50 μg/ml ampicillin and 25 μg/ml kanamycin until OD_600_ = 0.7 was reached. Then, expression was induced by adding 0.1% w/v L-arabinose, and cells were harvested after 2 h. Cells were disrupted using B-PER Bacterial Protein Extraction Reagent (ThermoFisher cat#78248) containing 6 mg/ml lysozyme. The GsSir2/Ago-NA complex was purified as described above, instead, all buffer solutions contained 100 mM NaCl.

To extract nucleic acids co-purified with the GsSir2/Ago complex, 800 μL of Roti-phenol/chloroform/isoamyl alcohol (Carl-Roth cat#A156) was added to the 800 μL of purified protein-NA fractions in 5PRIME Phase Lock Gel tubes (Quantabio cat#733-2477). The upper aqueous phase was isolated and 0.1 volumes of 1 M sodium acetate, 3 volumes of 100% ethanol and 10 μL glycogen (ThermoFisher cat#R0561) were added. This mixture was vortexed briefly and incubated at −20 °C for 20 hours. Samples were centrifuged for 20 min and the supernatant was removed from the pellet. The pellet was washed with cold (−20 °C) 70% ethanol. The pellets containing the copurified nucleic acids were dried for 20 min at room temperature, and pellets were resuspended in 30 μL water (free of nucleases).

Co-purified nucleic acids were dephosphorylated with FastAP Thermosensitive Alkaline Phosphatase (ThermoFisher cat# EF0651) and [γ-32P]-ATP (PerkinElmer) labelled with T4 polynucleotide kinase (PNK) (ThermoFisher cat#EK0031). Labelled nucleic acids were incubated with nucleases (ThermoFisher DNase I cat#18047019, RNase A/T1 cat# EN0551) for 30 min at 37 °C. After nuclease treatment, samples were mixed with RNA Gel Loading Dye (ThermoFisher cat# R0641), heated for 5 min at 95 °C and resolved on 20% denaturing (8 M Urea) polyacrylamide gels. The molecular weight marker used for RNA size identification was Decade Marker System (Ambion cat#AM7778) and 22 nt long RNA oligonucleotide. Radioactivity was captured from gels using phosphor screens and imaged using a Typhoon FLA 7000 laser-scanner (GE Healthcare).

In a control sample, a total RNA from induced cells was extracted using SPLIT RNA Extraction Kit (Lexogen cat#008). Then rRNA was removed using RiboCop for Gram-negative Bacteria (Lexogen cat#126).

### RNA sequencing and analysis

Half of extracted RNA was treated with T4 PNK (ThermoFisher cat#EK0031) according to the protocol of the manufacturer. Then T4 PNK treated and untreated RNA samples were converted to DNA libraries using Small RNA-Seq Library Prep Kit (Lexogen cat#052). Concentration and quality of libraries were measured with Qubit Fluorometer (ThermoFisher) and 2100 Bioanalyzer (Agilent).

Both libraries were sequenced using Illumina MiniSeq sequencing with single-end reads and 75 bp read length. Single-end reads were processed by trimming adapters with AdapterRemoval v2.3.0^43^. Then the processed reads were aligned to the *E. coli* str. K12 substrain DH10B genome (GenBank: CP000948.1) and the additional pBAD/HisA_TwinStrep_TEV_GsSir2/Ago, pCDF_Kn plasmids (Supplementary Table 1) using BWA-MEM v0.7.17^44^. In order not to filter out shorter reads during the alignment process, aligned reads with MAPQ values greater or equal to 15 were chosen. FastQC v0.11.8^45^ was used for read quality control and SAMtools v1.7^46^ – for indexing, sorting and analysing alignment files. A custom script (fragmentation-bias.jl) in combination with Weblogo v3.7.4^47^ were used to produce nucleotide frequency plots. The custom script had to be implemented to ensure that only aligned reads would be used for nucleotide frequency analysis. Gene enrichment analysis was performed with bedtools v2.26.0^48^ and FPKM_count.py v4.0.0 of RSeqQC package^49^. IGV v2.5.2^50^ was mainly used to inspect and visualise read coverage along the genomes. Control DNA library of a total RNA was prepared using CORALL Total RNA-Seq Library Prep Kit (Lexogen cat#095). Concentration and quality of the library was measured with Qubit Fluorometer (ThermoFisher) and 2100 Bioanalyzer (Agilent) according to the protocol of the manufacturer. Control DNA library was sequenced using Illumina NextSeq sequencing with pair-end reads and 75 bp read length. The read processing, alignment and alignment analysis were analogous to those samples from Illumina MiniSeq sequencing.

### Preparation of *E. coli* cells for NAD^+^ quantification

Overnight cultures of single colonies of *E. coli* DH10B strain harbouring a pBAD-His construct with either wt GsSir2/Ago or mutant system (GsSir2/Ago-HSH or GsSir2(D230A)/Ago), or empty vector (negative control) were diluted and grown in LB broth (BD) supplemented with respective antibiotics (50 μg/ml ampicillin and 25 μg/ml streptomycin) at 37 °C until they reached OD_600_ of 0.4–0.5. Cell cultures were either induced to express the protein or not (control samples). L-Ara (0.1% final concentration) was added to induce protein expression. Induced and non-induced cultures were harvested 2 hours later. The cultures were normalized to OD_600_ of approximately 0.7 and the pellet from 1 mL of culture suspension was stored at −80 °C until further analysis. All cell pellets were lysed by adding B-PER solution (Thermofisher Scientific) supplemented with 6 mg/ml lysozyme (62971, Fluka) for 20 min at room temperature while gently rocking (Multi Bio 3D Mini-Shaker, Biosan). Cell debris was removed by centrifugation and metabolites were isolated by phenol:chloroform:isoamyl alcohol (PCI) (25:24:1, v/v/v) extraction. Metabolites were stored at −20 °C until MS-HPLC analysis. Additionally, the endogenous NAD^+^ concentration was estimated using NAD/NADH Quantitation Kit (Sigma Aldrich, cat# MAK037) from four independent measurements.

### *In vitro* NADase assay

Reaction mixtures with a volume of 25 μL were prepared with the following final concentrations: 0.5 μM GsSir2/Ago or mutant complex, 50 μM NAD^+^, 1× Tango buffer (33 mM Tris-acetate (pH 7.9 at 37 °C), 10 mM magnesium acetate, 66 mM potassium acetate, 0.1 mg/ml BSA, ThermoScientific, cat#BY5), 1 mM DTT, 0.5 μM 5’P-RNA guide (TF-A) and/or 0.5 μM ssDNA (MZ-949 or MZ-589) (Supplementary Table 2). Reactions with RNA guide were preincubated for 15 min at 37 °C, then ssDNA was added and the mixture was incubated for 1 h at 37 °C. 3 μL of each sample was used as input for the NAD/NADH Quantitation Kit (Sigma-Aldrich, cat# MAK037) according to the instructions provided by the manufacturer. All experiments were performed in triplicates. These samples were also used for mass spectrometry.

### Mass spectrometry of NAD^+^

To quantitate NAD^+^ bound to GsSir2/Ago and CcSir2/Ago complexes high-performance liquid chromatography-mass spectrometry/mass spectrometry (HPLC-MS/MS) analysis was used. First, purified pAgos complexes were diluted to 5 μM in a buffer containing 20 mM Tris-HCl (pH 8.0 at 25 °C), 200 mM NaCl. Then 20 μl of the solution was incubated at 70 °C for 20 min and centrifuged for 30 min (16,100 g at 4 °C) to remove unfolded proteins. The supernatants and NAD^+^ standards were analysed by Electrospray Ionization mass spectrometry (ESI-MS) using an integrated HPLC/ESI-MS system (1290 Infinity, Agilent Technologies/Triple Quadrupole 6410, Agilent Technologies), equipped with a Supelco Discovery®HS C18 column (7.5 cm × 2.1 mm, 3 μm), Agilent Technologies. HPLC/ESI-MS/MS was performed using two ion transitions to detect NAD^+^ in the samples: 662.1→540.1 and 662.1→426.0. Ion transition 662.1→540.1, as it is the most abundant, was used for the quantitative analysis. Mobile phase A was 5 mM ammonium acetate in water, pH 7.0 and mobile phase B was 5 mM ammonium acetate in methanol, pH 7.0. The HPLC parameters were as follows: flow 0.25 ml/min; column temperature 30°C; 0-3 min, 0%B; 3-9 min, 0-40%B; 9-10 min, 40-100%B; 10-13 min, 100%B. The MS was operated using negative electrospray ionisation at 2500 V, the gas temperature was set to 300 °C, the fragmentor voltage was 135 V. Multiple reaction monitoring (MRM) was used with a collision energy of 15 V to measure ion m/z 540.1 (ion transition 662.1→540.1) and also with a collision energy of 20 V to measure ion m/z 426.0) (ion transition 662.1→426.0).

To quantitate endogenous NAD^+^ HPLC-MS analysis was performed by Electrospray Ionization mass spectrometry (ESI-MS) using an integrated HPLC/ESI-MS system (1290 Infinity, Agilent Technologies/Q-TOF 6520, Agilent Technologies), equipped with a Supelco Discovery®HS C18 column (7.5 cm × 2.1 mm, 3 μm), Agilent Technologies. The samples were investigated in both negative and positive ionization modes. For negative ionization mode, solvents A (5 mM ammonium acetate in water pH 7.0) and B (5 mM ammonium acetate in methanol, pH 7.0) were used. For positive ionization mode, solvents C (0,02% formic acid in water) and D (0,02% formic acid in acetonitrile) were used. In both cases elution was performed with a linear gradient of solvents at a flow rate of 0.3 ml/min at 30 °C as follows: 0–5 min, 0% B; 5–18 min, 20% B; 18–22 min, 100% B, 22–27 min 100% B. Ionization capillary voltage was set to 2500 V and fragmentor to 150V. A list of compounds that could be expected to be products of NAD^+^ hydrolysis and relative m/z value is as follows: ADPR [M-H]- m/z=558.0644, cADPR [M-H]- m/z =540.0538, AMP [M-H]- m/z=346.0558, cAMP [M-H]- m/z=328.0452, ADP [M-H]- m/z=426.0221, cADP [M-H]- m/z=408.0116, nicotinamide [M+H]+ m/z= 123.0553, adenine [M+H]+ m/z=136.0618. Only traces of AMP and ADP were detected in all samples; other products from the list were absent.

NAD^+^ hydrolysis products generated by GsSir2/Ago *in vitro* were analysed as above. Using negative ionization mode only accumulation of ADPR was detected.

## Data availability

All data are available in the paper and the supplementary material. In addition, small and total RNA sequencing data are available on the NCBI Sequence Read Archive under BioProject ID PRJNA851009. SAXS data are available in the Small Angle Scattering Biological Data Bank SASBDB under SASBDB ID SASDNH2: https://www.sasbdb.org/project/1486/cuudir1lvf/. Plasmid sequences used in this work are available at https://www.benchling.com, with exact links for each plasmid provided in Supplementary Table 1.

*Geobacter sulfurreducens, Caballeronia cordobensis*, and *Paraburkholderia graminis* genomes (respective GenBank accessions: GCA_000210155.1, GCA_001544575.2 and GCA_000172415.1) and all associated sequence and annotation data were obtained from NCBI (ftp://ftp.ncbi.nlm.nih.gov/genomes/Bacteria/). Searches through Pfam (http://pfam.xfam.org/), SwissProt (https://www.expasy.org/resources/uniprotkb-swiss-prot), and PDB (https://www.rcsb.org/) databases were performed. PDB structures mentioned in this study: 5AWH, 4N41, 5UX0, 6LHX, 2H4F.

## Code availability

The Julia script used to identify nucleotide frequency in the beginning of the aligned reads and prepare input for Weblogo program is available at GitHub repository: https://github.com/agrybauskas/argonaute-bound-rna-manuscript

## Acknowledgements

We thank the Siksnys laboratory members for their comments on the manuscript and fruitful discussion. This work was supported by European Social Fund [09.3.3-LMT-K-712-01-0126 to V.S.] under a grant agreement with the Research Council of Lithuania (LMTLT), the Israel Science Foundation [grant ISF 296/21 to R.S.], and the Deutsche Forschungsgemeinschaft [SPP 2330, grant 464312965]. The authors thank Tomás de Garay at Refeyn Ltd. for the mass photometry experiments. Funding for open access charge: Vilnius University.

## Author Contributions

V.S. and M.Z. designed the study; K.T. and Č.V. performed bioinformatics and structural modelling; A.L. performed the phage restriction experiments; E.S., D.D., E.G., S.A., U.T. performed the plasmid transformation experiments; A.S. purified the proteins and performed the SEC-(MALS) experiments; R.G. performed the SEC experiments; E.M. performed the SAXS experiments; E.Z., E.G., D.D. and R.G. performed the EMSA experiments; E.Z. and E.J. reconstituted the GsSir2/Ago-RNA complex and performed the biochemical analysis; A.G. performed the RNA-seq analysis; A.R. performed the mass spectrometry analysis; D.D. performed the NAD^+^ determination experiments *in vivo*; E.Z. performed the NAD^+^ hydrolysis experiments *in vitro*; R.S., V.S. and M.Z. analysed the data; M.Z. wrote the initial manuscript with input from E.G., R.S., V.S. and other authors. All authors approved the final version.

## Notes

During the revision of this manuscript, a paper was published that shows plasmid-induced degradation of NAD^+^ *in vivo* by short pAgo-associated TIR-APAZ systems (named SPARTA)^52^. The paper also shows that expression of Sir2/Ago systems (named SPARSA) in *E. coli* triggers NAD depletion.

## Extended data figures

**Extended Data Fig. 1.**
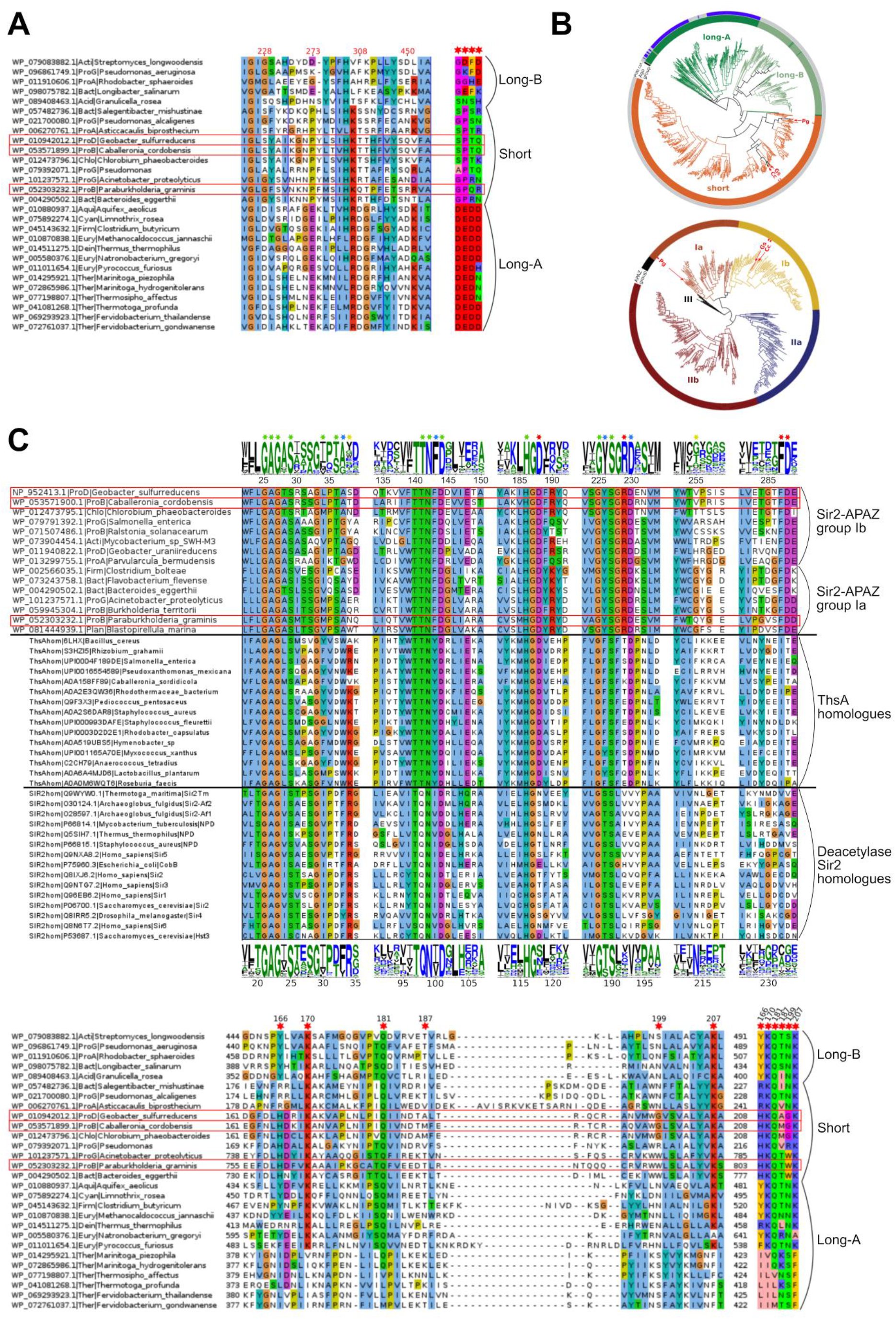
Bioinformatic analysis. **A**, PIWI catalytic tetrad DEDX alignment. The 4 catalytic residues (red numbers indicate positions of corresponding GsAgo positions) are shown in 4 motifs of ±3 positions. The motifs are separated by vertical blue lines. Sequence names consist of the following: NCBI sequence ID, abbreviated phylum (e.g., ‘ProG’ – gamma-proteobacteria) and organism name. **B**, Top - a circular phylogenetic tree was generated according to supplementary data provided with Ryazansky et al.^8^ Long-A pAgo variants are coloured in green (truncated variants without the PAZ domain, light green), long-B pAgo proteins are light green (truncated variants without PAZ, green), and short pAgo proteins are orange. pAgo proteins containing the catalytic tetrad DEDX in their PIWI domain are indicated in blue on the outer circle; pAgos with inactivated PIWI domain are indicated in light grey on the outer circle. pAgo proteins of the GsSir2/Ago, CcSir2/Ago and PgSir2-Ago systems are indicated by ‘Gs’, ‘Cc’ and ‘Pg’, respectively. Bottom - Circular phylogenetic tree of APAZ domains. The circular phylogenetic tree of the five groups of APAZ domains was generated using APAZ domain alignments from Ryazansky et al.^8^ supplementary file 7. **C**, Top - Combined alignment of Sir2 domains. Alignment consists of 3 parts, separated by horizontal black lines. In the top part, the Sir2 domain sequences of the GsSir2, CcSir2, PgSir2-Ago and homologues are shown. Logos above depict the conservation of Sir2 domains of Ia and Ib groups. The indicated position numbers correspond to the GsSir2 sequence. In the bottom part, homologues (sirtuins) of catalytically active Thermotoga maritima Sir2 (TmSir2) deacetylase are shown. Logos below indicate the conservation of these homologues. The position numbers correspond to the TmSir2 sequence. Sequences of six motifs that include all positions that form the NAD^+^-binding pocket, as seen in the TmSir2 structure (PDB ID 2H4F) are shown. Sequence names for the top alignment consist of sequence ID, abbreviated phylum and organism name. Sequence names for bottom alignment all start with “Sir2hom” followed by sequence ID, organism name and short protein name (based on annotation). Stars above the logos indicate residues in the NAD^+^-binding pocket of canonical sirtuins (e.g., TmSir2) that are also conserved. Star colours indicate conservation between the two groups: green – conserved in both canonical sirtuins and GsSir2-like; blue – conserved in both groups, but different; yellow – conserved only in canonical sirtuins; red – conserved only in GsSir2-like proteins. In the middle, alignment of ThsA homologues with Sir2 domains. Bottom - MID domain alignment. Red stars indicate positions of amino acids involved in the binding of the 5’-P end of the guide nucleic acid. The numbering above corresponds to the GsAgo sequence. Additionally, concatenated alignment of just the 6 indicated positions is shown on the right. The three sequences of interest are indicated with red rectangles. Numbers on the left and right of the alignment indicate the first and last positions in the alignment for each sequence.

**Extended Data Fig. 2.**
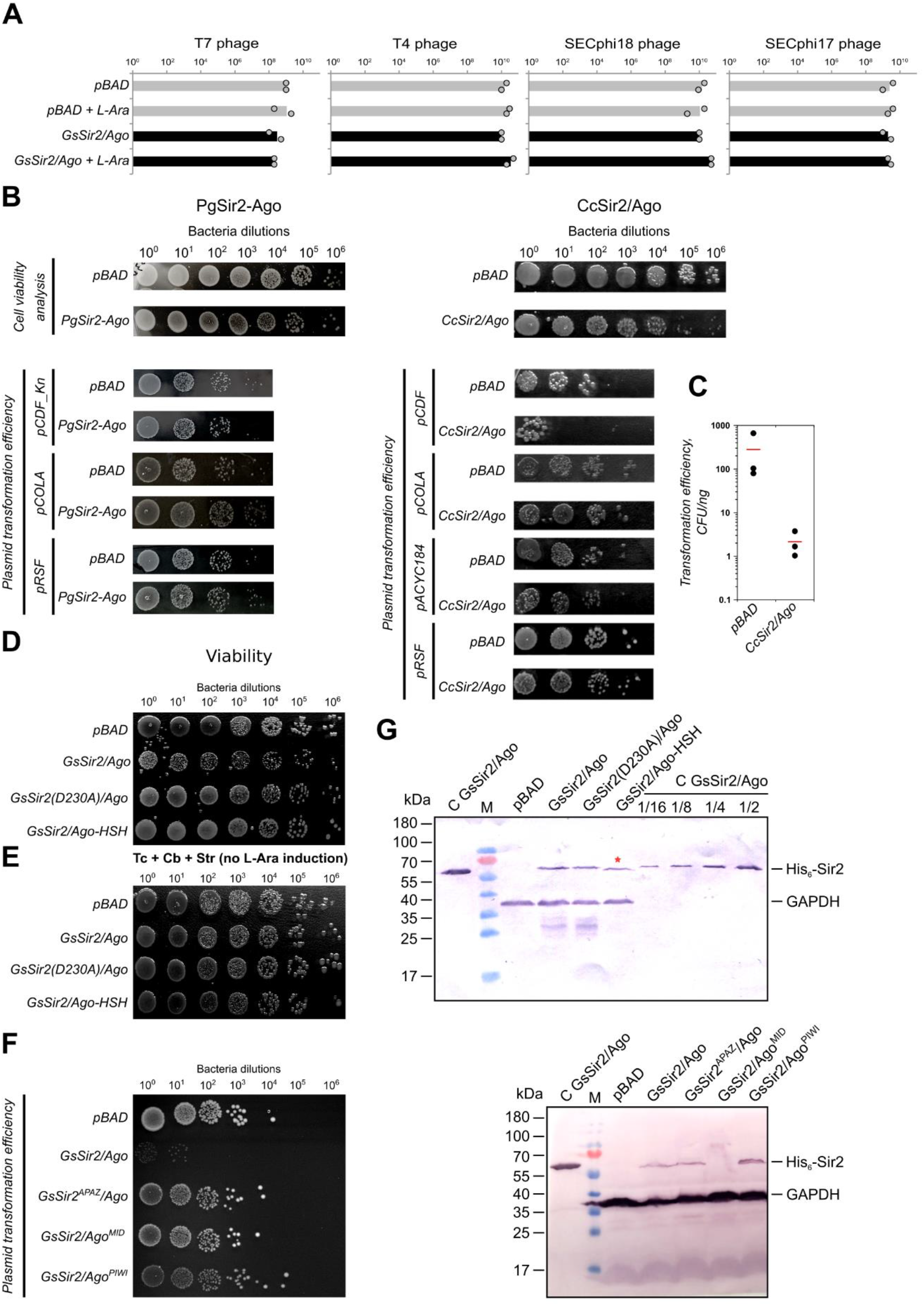
*In vivo* characterization of Sir2/Ago systems. **A,** Efficiency of plating (EOP) of 4 phages infecting *E. coli* cells with and without the GsSir2/Ago system from *Geobacter sulfurreducens*, where the GsSir2/Ago system exhibits no defence activity. The x-axis represents the number of p.f.u. Shown are the means of two replicates in the absence and in the presence of the inducer L-arabinose (L-Ara), with individual datapoints superimposed. Grey bars represent efficiency of plating (EOP) on pAgo-lacking cells and black bars are EOP in pAgo-containing cells. **B**, Left - qualitative characterization of plasmid restriction capabilities of PgSir2-Ago system in *E. coli* strain DH10B. Top: comparison of cell viability in the presence or absence of plasmid-borne PgSir2-Ago expression. Bottom: comparison of plasmid transformation efficiencies in the presence or absence of plasmid-borne PgSir2-Ago expression. Right - qualitative characterization of plasmid restriction capabilities of CcSir2/Ago system in *E. coli* strain BL20-AI: top - comparison of cell viability in the presence or absence of plasmid-borne CcSir2/Ago expression. Bottom - comparison of plasmid transformation efficiencies in the presence or absence of plasmid-borne CcSir2/Ago expression. **C**, Quantification of transformation efficiencies for pCDF plasmid with CcSir2/Ago system (three independent replicates, the red line represents average transformation efficiency). **D**, Cell viability control of BL21-AI, expressing GsSir2/Ago and mutants. **E**, Control for Fig. 2E – cells contain the pCDF plasmid, however, expression of the GsSir2/Ago system is not induced. **F,** Qualitative evaluation of pCDF plasmid transformation efficiency in *E. coli* cells carrying GsSir2/Ago mutants (GsSir2^APAZ^/Ago, GsSir2/Ago^MID^ and GsSir2/Ago^PIWI^) of the putative surface of the interaction with nucleic acids. **G**, Expression analysis of GsSir2/Ago assayed by Western blot. Top: semiquantitative Western blot of the wt GsSir2/Ago complex and its mutants. Numbers above the lanes indicate which part of control protein amount is loaded. The red star shows the lane where the His-tag is on the C-terminus of Ago, rather than the N-terminus of Sir2. GAPDH, loading control. Bottom: Expression analysis of GsSir2/Ago mutants of the putative surface of the interaction with nucleic acids. Three replicates.

**Extended Data Fig. 3.**
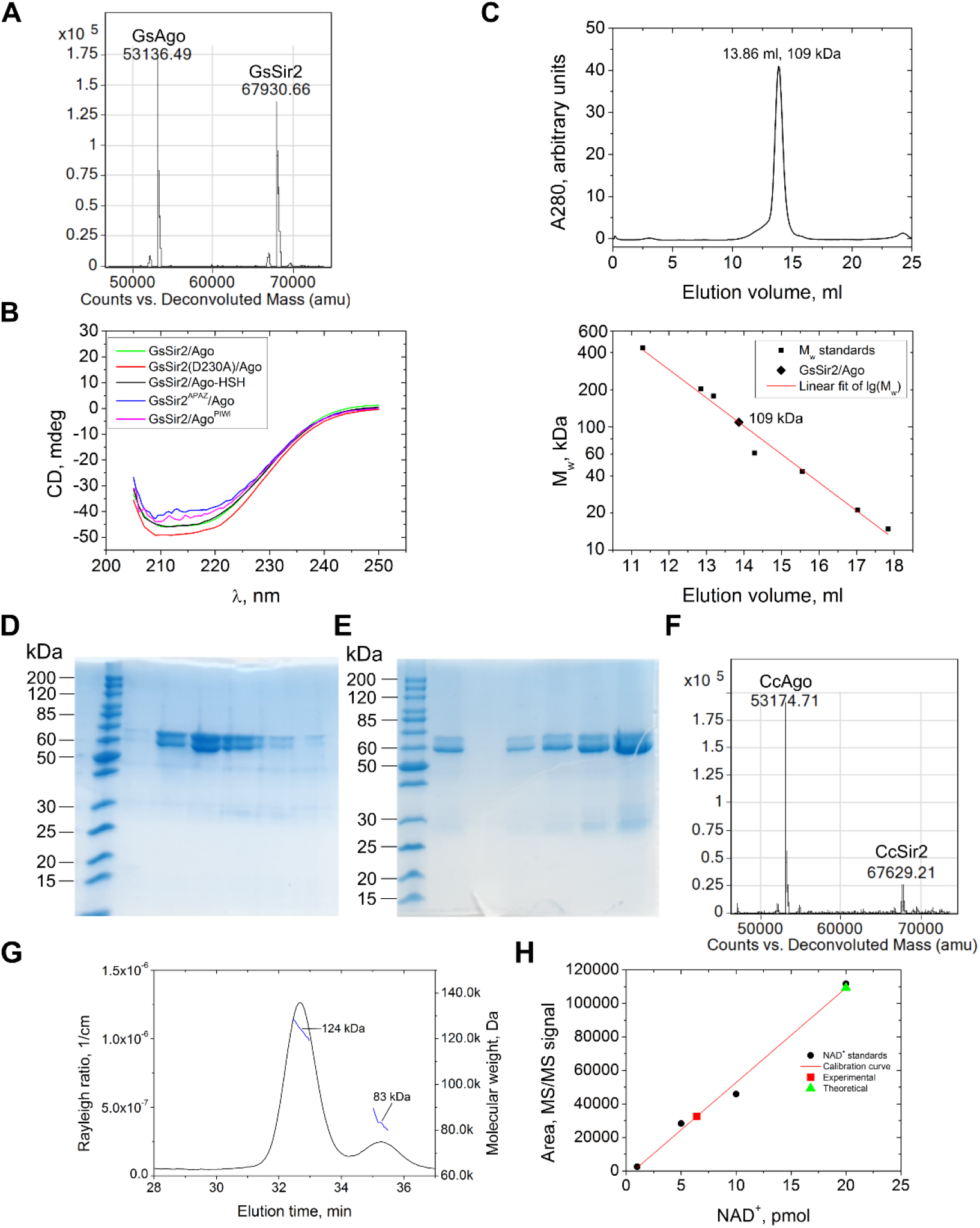
Purification and characterization of Sir2/Ago complexes. **A**, MS analysis of wt GsSir2/Ago complex. The theoretical Mw of the GsSir2 and GsAgo proteins (without 1st Met) are 67929.51 Da and 53135.61 Da, respectively. **B**, CD spectra of wt GsSir2/Ago and mutants. Mutant spectra are similar to that of a natively folded protein. **C**, Size-exclusion chromatography of wt GsSir2/Ago, showing elution volume and comparing to mass standards. According to mass spectrometry of the purified GsSir2/Ago complex, the molar mass of the GsSir2/Ago heterodimer is 121 kDa. **D**, SDS-PAGE analysis of fractions containing the CcSir2/Ago complex eluted from Heparin column. Densitometric inspection shows that Sir2 and Ago proteins are in the ratio ~1:1. Single replicate. **E**, SDS-PAGE analysis of the CcSir2/Ago stock after dialysis against a storage buffer. Various amounts of the stock solution were loaded on the gel. Densitometric inspection shows that Sir2 and Ago proteins are in the ratio ~0.3:1. Single replicate. **F**, MS analysis of the CcSir2/Ago complex. The experimental masses (53174.71 Da and 67629.21 Da) are close to the theoretical molecular masses of the Ago protein (53173.91 Da) and the Sir2 protein with the truncated tag at the N terminus (67626.24 Da). **G**, SEC-MALS analysis of the CcSir2/Ago complex. The experimental mass of 124 kDa is close to the theoretical molecular mass of the CcSir2/Ago heterodimer (121 kDa). **H**, MS/MS calibration curve of NAD^+^ standard (marked in black) and the observed amount of NAD^+^ (marked in red) in the CcSir2/Ago complex (20 pmol according to the Ago protein). The discrepancy between the expected amount of NAD^+^ (20 pmol, marked in green) and the actual amount (6.45 pmol, marked in red) was due to the decrease of the Sir2 protein in the CcSir2/Ago preparation (see **E**).

**Extended Data Fig. 4.**
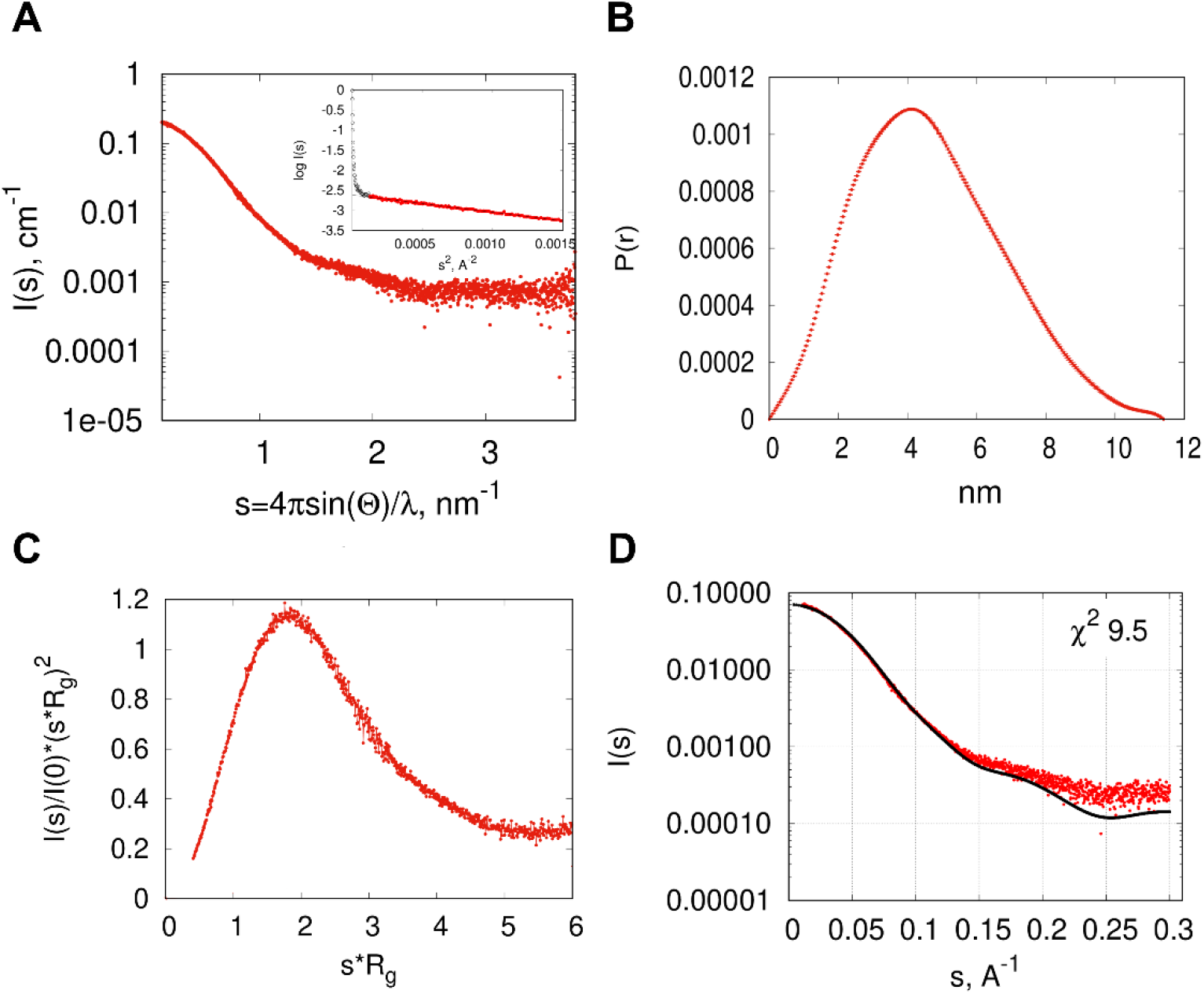
SAXS data. **A**, Scattering data on an absolute scale. Linear Guinier plot of the initial part of the scattering curve is in the insert. Points cut from the further processing are shown with empty black symbols. **B**, Kratky plot, normalized by Rg and I(0) parameters. **C**, Pair distance distribution function. **D**, CRYSOL Fit of the scattering curve calculated from the GsSir2/Ago AlphaFold model (black curve) with SAXS data (red points).

**Extended Data Fig. 5.**
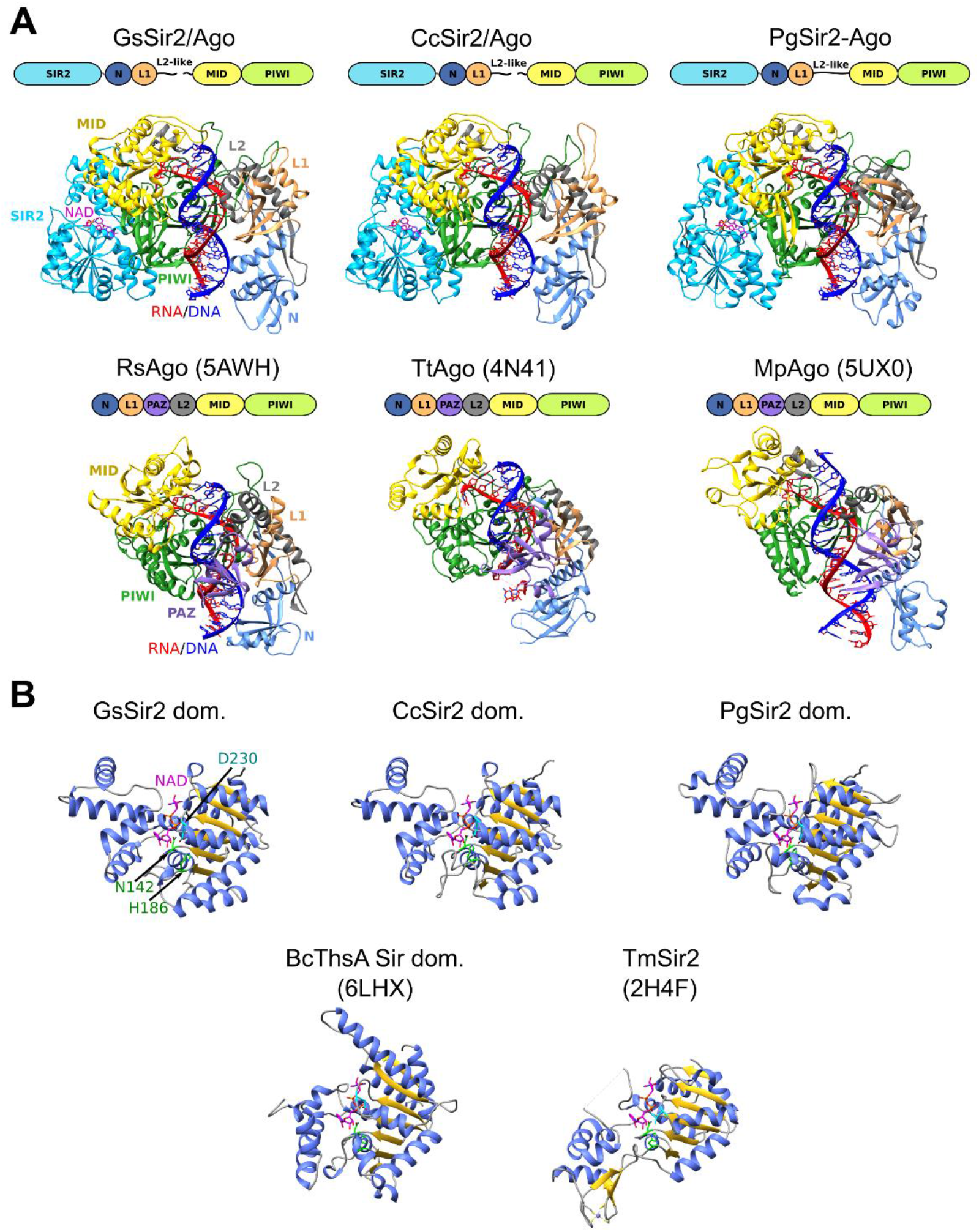
Structural analysis. **A**, Comparison of GsSir2/Ago, CcSir2/Ago and PgSir2-Ago AlphaFold models with the X-ray structures of long pAgos. Structures are coloured by domains, schematic domain architecture is given above each structure. Guide RNA and target DNA strands are coloured red and blue, respectively. PDB ID codes for long pAgo structures are given in parentheses. RsAgo represents long-B group, TtAgo – longA group of long pAgos (based on Ryazansky et al. classification^8^). MpAgo has a distinct OH-type MID domain that is specific to 5’-OH instead of phosphate (MID domain classification – Ryazansky et al.^8^, MpAgo MID domain biochemical assay - Kaya et al.^51^). In Sir2/Ago models the N, L1 and L2-like domains previously identified as the APAZ domain correspond to the analogous domains of long pAgos. **B**, Gs, Cc and Pg Sir2 structural models (cut from full-length models) compared to canonical Sir2 deacetylase TmSir2 and the Sir2 domain of Thoeris defence system protein ThsA. Structures are coloured based on secondary structure. Positions corresponding to ThsA Sir2 N112 and H152 are indicated in green. These residues have been shown to be critical for NAD^+^ hydrolysis in ThsA^24^. GsSir2 D230 and corresponding positions in other structures are indicated in cyan. NAD^+^ was also superimposed on the ThsA Sir2 structure from TmSir2.

**Extended Data Fig. 6.**
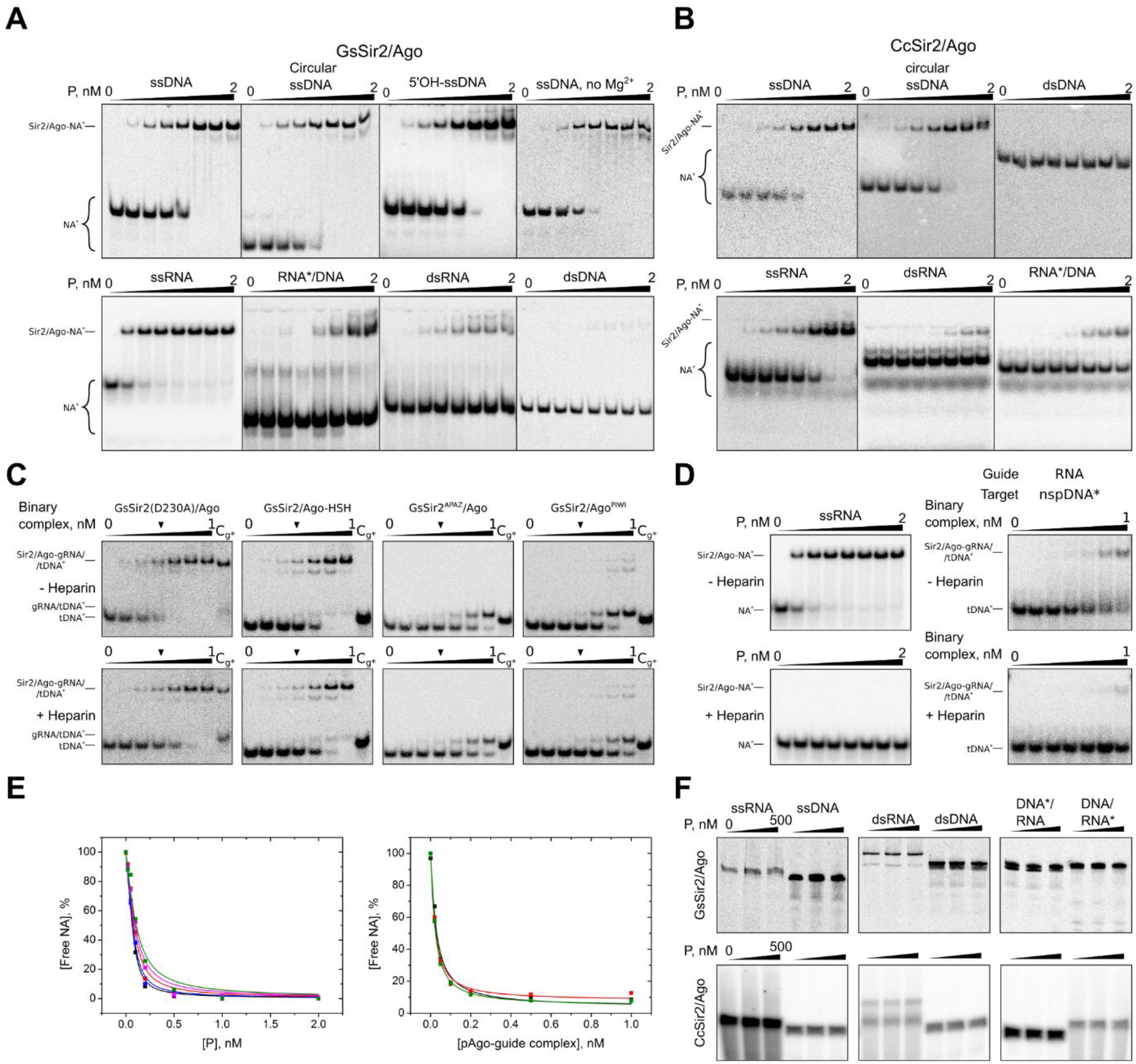
EMSA and nucleic acid cleavage experiments. **A-B**, Binding of single- and double-stranded oligonucleotides by wt GsSir2/Ago and wt CcSir2/Ago, respectively. A radiolabelled strand indicated by the asterisk. **C**, Binding of complementary DNA targets by GsSir2/Ago binary complexes pre-loaded with RNA guide containing 5’-phosphate terminus in the presence or absence of heparin. To show that no displacement of the radiolabelled guide by the target strand is observed, a control (Cg*) equivalent to the experimental lane marked by a black triangle, but with the guide, rather than the target, bearing the radioactive label, was performed. **D**, Control EMSA experiments of ssRNA guide binding by wt GsSir2/Ago (left) and non-complementary DNA target binding by the binary wt GsSir2/Ago - gRNA complex in the presence and absence of heparin. **E**, Representative binding fit curves of several independent replicates used to calculate Kd of ssRNA guide binding by wt GsSir2/Ago (left) and of target DNA binding by wt GsSir2/Ago-gRNA complex (right). **F**, (No) cleavage activity of various DNA and RNA oligonucleotides by wt GsSir2/Ago and wt CcSir2/Ago. Reaction products were resolved on a 21% denaturing polyacrylamide gel. In heteroduplexes, the asterisk indicates the radiolabelled strand. For panels **A-D** and **F**, at least three independent replicates were performed for each experiment.

**Extended Data Fig. 7.**
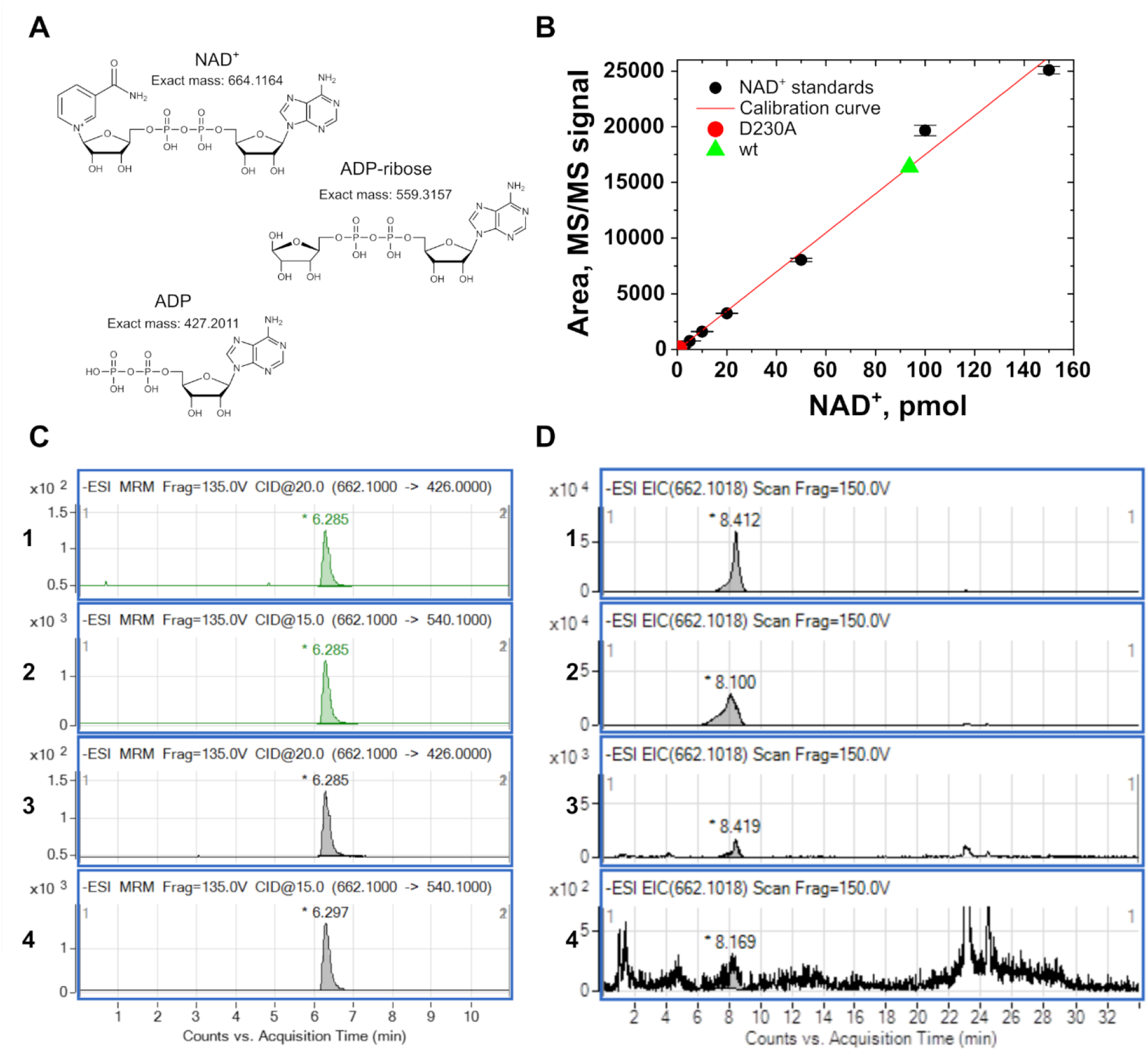
The GsSir2/Ago complex binds NAD^+^ and causes its depletion. **A**, Two ion transitions were used to detect NAD^+^ in the analysed samples: 662.1→540.1 and 662.1→426.0. **B**, MS/MS calibration curve of NAD^+^ standard (marked in black, two replicates) and the observed amount of NAD^+^ in two samples: 93.7 pmol in the wt GsSir2/Ago sample (marked in green), 0.7 pmol in the sample D230A (marked in red). Black dots represent the means of two replicates and error bars are the standard deviation. **C**, Detection of NAD^+^. Comparison of the extracted LC-MS/MS chromatograms: ion transition 662.1→426.0 of wt GsSir2/Ago sample (panel 1) and NAD^+^ standard (panel 3); ion transition 662.1→540.1 of wt GsSir2/Ago sample (panel 2) and NAD^+^ standard (panel 4). Green curves - wt GsSir2/Ago sample, grey curves – NAD^+^ standard. **D**, Mass chromatogram. For NAD^+^ detection, an extracted ion current (EIC) for [M-H]- m/z = 662.1018 was used. The comparison of EIC signals shows that the amount of NAD^+^ in the samples of the non-induced GsSir2/Ago system in the absence (panel 1) and presence (panel 2) of pCDF plasmid is almost the same, while a significant decrease is observed in the sample of the induced GsSir2/Ago system in the absence of pCDF plasmid (panel 3) and only traces of NAD^+^ are detected in the presence of pCDF plasmid (panel 4).

## Supplementary Information

### Supplementary Note 1

#### Bioinformatic analysis of the selected Sir2/Ago systems

Closer inspection of the genomic neighbourhood of the selected pAgos showed that no other conserved operons are formed with the pAgo genes. However, the observed enrichment of putative restriction endonucleases, mobile genetic elements (e.g. transposases, integrases) and toxin-antitoxin systems in the neighbourhood of the GsAgo and CcAgo genes indicates that they could be a part of so-called bacterial defence islands^1^. All three systems have MID-PIWI domains, characteristic of Agos, but their PIWI domains are catalytically inactive (Fig. 1A). Importantly, this derived feature is shared with the evolutionary closest group of long pAgos (Extended Data Fig. 1C). Although overall sequence similarity between the MID-PIWI regions of selected systems (Gs, Cc and Pg) and long pAgos is rather low (~15-20% sequence identity), their MID domains do have conserved residues associated with binding of the 5’-end of the guide strand^2^ (Extended Data Fig. 1E). The conserved motif of MID domains of all three systems is of the HK-type, which is more similar to the 5’-P-end binding motif present in long pAgos such as RsAgo or TtAgo than to the 5’-OH-end binding motif found, for example, in MpAgo^2^. Unlike long pAgos, short pAgos lack the PAZ domain and are rather associated with an APAZ (analogue of PAZ) domain (Fig. 1A)^3^. By phylogeny of the APAZ domain region, PgSir2-Ago belongs to Ia, and GsSir2/Ago and CcSir2/Ago to Ib groups, respectively (Extended Data Fig. 1B)^2^. The sequences of Sir2 domains associated with the three studied pAgos are similar to canonical sirtuins^3^, and also have an identifiable signature of a putative NAD^+^-binding pocket (Extended Data Fig. 1C). Yet there are many differences from canonical sirtuins, and the identified conserved amino acid positions found only in pAgo-associated Sir2 domains suggest that these domains might have a function different from the typical deacetylase activity of sirtuins (Extended Data Fig. 1C). pAgos-associated Sir2 domains also show similarities to NADase domains of ThsA proteins from the Thoeris anti-phage systems (Extended Data Fig. 1C)^4,5^.

#### GsSir2/Ago complex resembles long PAZ-free pAgos containing an additional effector domain

To get an insight into the 3D structure of GsSir2/Ago and the other two systems, we used AlphaFold^6^ to generate corresponding structural models. Based on high AlphaFold confidence values (plDDT>87; pTM>0.82) and favourable VoroMQA^7^ statistical energy scores (>0.53) the models for all three systems might be expected to be of comparable quality with experimental structures. Consistent with clear homology between the three systems the corresponding models showed close structural similarity with each other (Extended Data Fig. 5). Therefore, we focus only on the GsSir2/Ago complex. In this complex, the Sir2 and Ago subunits bind together to form a structure similar to that of a single-chain long pAgo protein such as RsAgo (Extended Data Fig. 5). Based just on a structural similarity search against PDB, GsSir2/Ago appears to be most similar to the TtAgo structure classified as a long-A pAgo^2^. Another close structural match is RsAgo, a member of long-B pAgos^2^. GsAgo shares 17% identical residues with both TtAgo and RsAgo. However, since RsAgo, just like short pAgo proteins, has a catalytically inactive PIWI domain, a comparison of GsSir2/Ago with RsAgo might be biologically more relevant.

The APAZ part of the GsSir2 chain is structurally similar to the combination of N, L1 and L2 domains of RsAgo (similarity to the N domain has been already proposed earlier^8,9^). However, the L2 linker domain in GsSir2/Ago corresponds to the two fragments, C-terminus of Sir2 and N-terminus of Ago. Importantly, the PAZ domain, required for the 3’-end recognition of the guide strand in long pAgos, is missing from the GsSir2/Ago structure altogether. Additionally, GsSir2/Ago has a smaller N domain than long pAgos, which could also alter nucleic acid binding. PIWI and MID domains are structurally similar to corresponding RsAgo domains. Notably, GsSir2/Ago has a longer loop (residues 268-272) compared to the corresponding region in RsAgo. This loop shows some steric overlap with the copied-in RNA/DNA duplex, but presumably, its conformation may adjust to accommodate RNA/DNA heteroduplex. The loop contains two positively charged residues (R269, K270) that might be involved in the binding of the nucleic acid backbone.

In the structural model, the N-terminal Sir2 domain of the GsSir2 protein is attached to the C-terminal domain corresponding to the N-terminal region of a long pAgo through a long linker. In effect, the two domains of GsSir2 are positioned at the opposite extremes of the complex (Sir2 domain is bound to the PIWI domain whereas the C-terminal domain is bound to the MID domain). Sir2 domain of GsSir2/Ago is structurally similar to canonical Sir2 proteins suggesting it could bind NAD^+^ in a similar way. However, the overall structurally closest homolog is the Sir2 domain of the ThsA protein from the Thoeris defence system^5^ (Extended Data Fig. 5). The ThsA residues, N112 and H152, shown to be essential for NAD^+^ hydrolysis have their counterparts in GsSir2 (N142 and H186). Interestingly, the loop containing H186 in different GsSir2/Ago models displays some conformational heterogeneity hinting at possible flexibility, which might be relevant for the activity regulation. Conservation of these two positions is also observed in other Sir2 homologs (Extended Data Fig. 1C). On the other hand, the GsSir2 D230 residue shown here to be important for NAD^+^ hydrolysis is conserved in ThsA Sir2 domain, but not in the canonical Sir2 proteins (e.g., V193 in TmSir2). In the AF2 structural model, the NAD^+^-bound Sir2 active site is hidden at the interface between the MID and PIWI domains. One may assume that binding of the complementary DNA target by the binary GsSir2/Ago-gRNA complex triggers a conformational change that opens and activates the Sir2 active site resulting in NAD^+^ hydrolysis.

#### The activated GsSir2/Ago depletes NAD^+^ enzymatically

Our results show that wt GsSir2/Ago tightly interacts with NAD^+^ (Fig. 5A, Extended Data Fig. 7). Therefore, there is a theoretical possibility that the NAD^+^ depletion in *E. coli* cells expressing the GsSir2/Ago system might be related to its tight binding by the activated GsSir2/Ago complex. However, this assumption can be ruled out based on our approximate calculations of relative levels of the GsSir2/Ago protein and NAD^+^ molecules in *E. coli* cells. According to our semiquantitative Western Blot analysis (Extended Data Fig. 2G), the GsSir2/Ago concentration (~1 μM) is more than ~1000-fold lower than that of NAD^+^ (~2.5 mM), which is in the range of the published NAD^+^ concentrations (~0.6-8 mM)^10^. Therefore, the activated GsSir2/Ago complex should act enzymatically to deplete such a large excess of NAD^+^.

### Supplementary methods

#### Sequences analysis

Lists of pAgo homologues and associated APAZ domains were retrieved from supplementary data of the Ryazansky et al. article^2^. *Thermotoga maritima* Sir2 homologues were collected from the SwissProt database^11^ using BLAST^12^ (1e-5 e-value cutoff). Full-length sequences were retrieved from NCBI. *Bacillus cereus* ThsA (PDB ID 6LHX) homologues were collected using BLAST from UniRef50^13^ database (1e-10 e-value cutoff). To remove other Sir2 homologues, these sequences were clustered with CLANS^14^. The cluster separated at p=1e-70 was selected as the representative ThsA group. Fragmented ThsA sequences and sequences missing one of the domains were discarded. Multiple sequence alignments were generated using MAFFT (l-INS-i mode for high accuracy)^15^. Jalview^16^ was used for multiple sequence alignment analysis, cutting and visualization. Sequence motif visualization was done using WebLogo 3 server^17^. Construction of combined multiple sequence alignment of GsSir2, TmSir2 and ThsA homologues was guided by the alignments between GsSir2, TmSir2 and BsThsA sequences obtained using the HHpred server^18^.

#### Phylogenetic tree construction

The phylogenetic trees were constructed with FastTree^19^ using the WAG model of amino acid substitution^20^ and the gamma model of rate heterogeneity. Prior to phylogenetic analysis, multiple sequence alignment positions containing more than 50% gaps were removed using trimAl^21^. Phylogenetic tree visualization was done with iTOL^22^.

#### Genomic neighbourhood analysis

For genomic neighbourhood analysis, the *Geobacter sulfurreducens*, *Caballeronia cordobensis*, and *Paraburkholderia graminis* genomes (respective GenBank accessions: GCA_000210155.1, GCA_001544575.2 and GCA_000172415.1) and all associated sequence and annotation data were obtained from NCBI (ftp://ftp.ncbi.nlm.nih.gov/genomes/Bacteria/). Genes in the neighbourhood of each pAgo (10 upstream and 10 downstream) were identified based on available genome annotations. To refine available functional annotations of these genes, searches through Pfam^23^ and PDB databases were performed using the HHpred server^18^.

#### Expression and purification of CcSir2/Ago complex

Expression vector constructs of the CcSir2/Ago system were used to transform *E. coli* BL21(DE3) strain. Transformed bacteria were grown at 37 °C in LB medium in the presence of 50 μg/ml ampicillin until OD_600_ = 0.7 was reached. Then, the medium was cooled to 16 °C temperature and proteins were expressed for 16 h by adding 0.1% w/v L-arabinose. Harvested cells were disrupted by sonication in buffer A (20 mM Tris–HCl (pH 8.0 at 25 °C), 1.0 M NaCl, 2 mM phenylmethylsulfonyl fluoride, 5 mM 2-mercaptoethanol), and cell debris was removed by centrifugation. TwinStrep-CcSir2/Ago complex was purified to > 90% homogeneity by chromatography through Strep-Tactin XT Superflow (Iba), HiLoad Superdex 200, HiTrap Heparin HP columns (GE Healthcare). Purified proteins were stored at −20 °C in a buffer containing 20 mM Tris-HCl (pH 8.0 at 25 °C), 500 mM NaCl, 2 mM DTT and 50% v/v glycerol. The identity of the purified proteins was confirmed by mass spectrometry. Protein concentrations were determined from OD_280_ measurements using the theoretical extinction coefficients calculated with the ProtParam tool available at http://web.expasy.org/protparam/. TwinStrep-CcSir2/Ago complex concentrations are expressed in terms of heterodimer.

#### CD analysis

To test whether the introduced mutations have an effect on secondary structures of the GsSir2/Ago, CD spectra of the proteins were recorded. Protein samples (5 μM of heterodimer) were prepared in 200 μl of the buffer (10 mM Tris-HCl (pH 8.0 at 25 °C), 50 mM NaCl). The circular dichroism spectra were recorded in triplicate with Jasco J-815 CD spectrometer (Jasco, Easton, MD) using a 1 mm quartz cuvette at a scanning speed of 50 nm/min from 200 to 250 nm. The temperature of samples was controlled at 25 °C using a Jasco temperature control device.

#### Construction and analysis of Sir2/Ago structural models

Structural models were generated using the AlphaFold method^6^ implemented as ColabFold^24^, an online Google Colaboratory notebook. For modelling, the ‘Alphafold2_advanced’ notebook (colab.research.google.com/github/sokrypton/ColabFold/blob/main/beta/AlphaFold2_advanced.ipynb) was used. GsSir2/Ago and CcSir2/Ago complexes were modelled as heterodimers, whereas the Pg sequence representing a fusion of Sir2 and a short pAgo (PgSir2-Ago) was modelled as a monomer. The modelling pipeline was run with default parameters except for the multiple sequence alignment (MSA) pairing, which was set to ‘paired+unpaired’. The best-of-five model in each case was selected using VoroMQA^7^. Structure similarity searches of models/domains against PDB were performed using Dali^25^. Structure analysis and visualization was performed using UCSF Chimera^26^. Putative binding sites of Sir2/Ago complexes were investigated by simply copying the RNA/DNA duplex and NAD^+^ from RsAgo (PDB ID: 5AWH) and TmSir2 (PDB ID: 2H4F) structures correspondingly after their superposition onto structural models. No attempts to remove possible clashes between protein models and either RNA/DNA or NAD^+^ were made.

#### Nucleic acid cleavage assay

The same linear oligonucleotides used for EMSA (Supplementary Table 2) were given as substrates for nucleic acid cleavage assay. To raise the total substrate concentration, 5’-^32^P radiolabelled oligonucleotides were mixed with appropriate cold 5’P-oligonucleotides at a ratio of 1:4 of hot:cold, and diluted to a working concentration of 100 nM in reaction buffer (33 mM Tris-acetate, pH 7.9, supplemented with 10 mM magnesium acetate, 66 mM potassium acetate, 0.1 mg/ml BSA, 5 mM DTT). Protein dilutions were made using the same reaction buffer to 2x final reaction concentration. Protein complexes and nucleic acids were mixed in final concentrations of 50 nM NAs and 0, 50 or 500 nM protein, incubated for 1 hour at 25 °C. The reaction was stopped with the addition of 2x 95% formamide dye and incubating for 5 min at 95 °C. Reaction products were resolved by denaturing PAA gel electrophoresis (21% 29:1 acrylamide/bis-acrylamide in TBE (89 mM Tris, 89 mM boric acid, 2 mM EDTA), supplemented with 8 M urea), visualised with a phosphor imager and analysed in OptiQuant software.

### Supplementary tables

**Supplementary Table 1.**
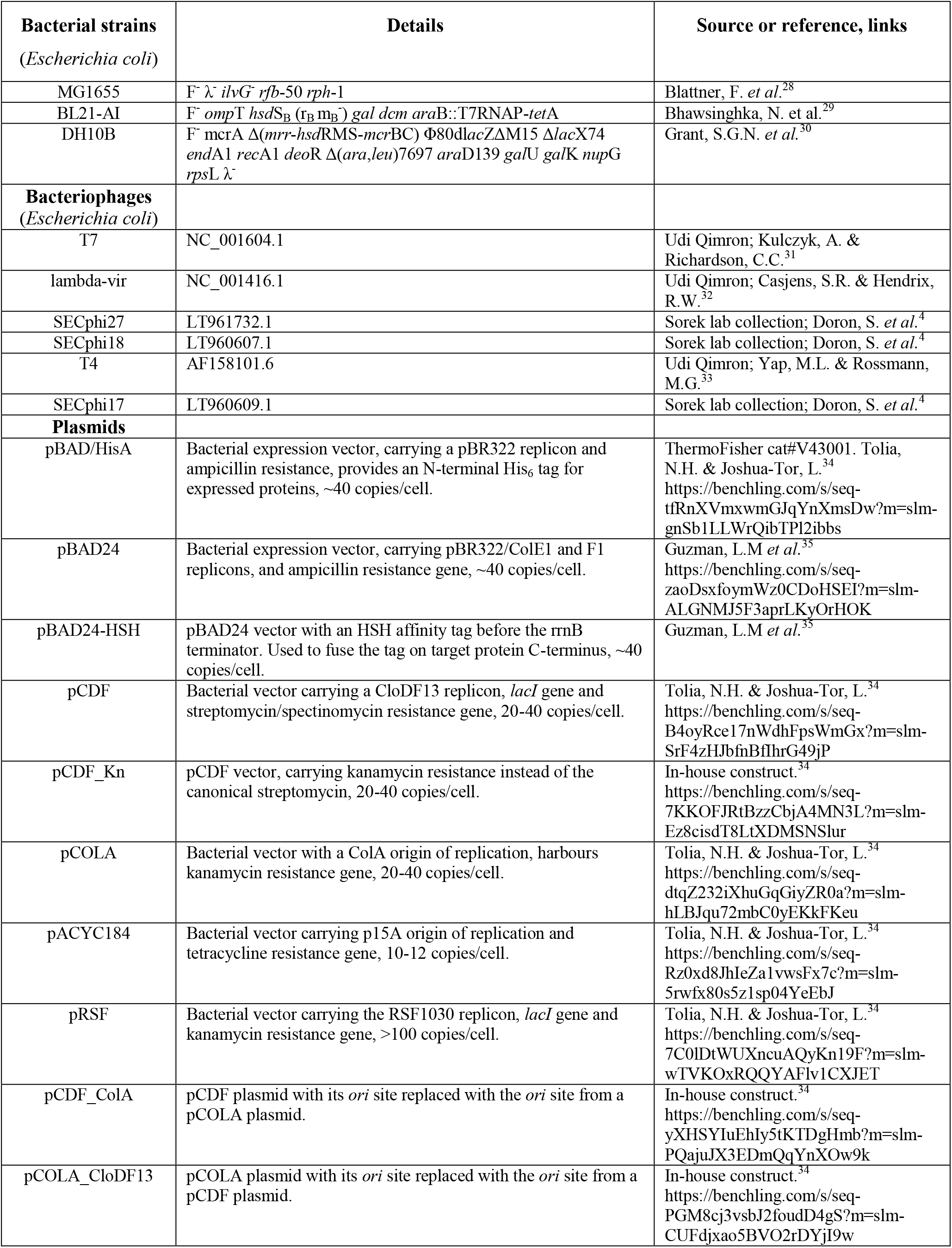
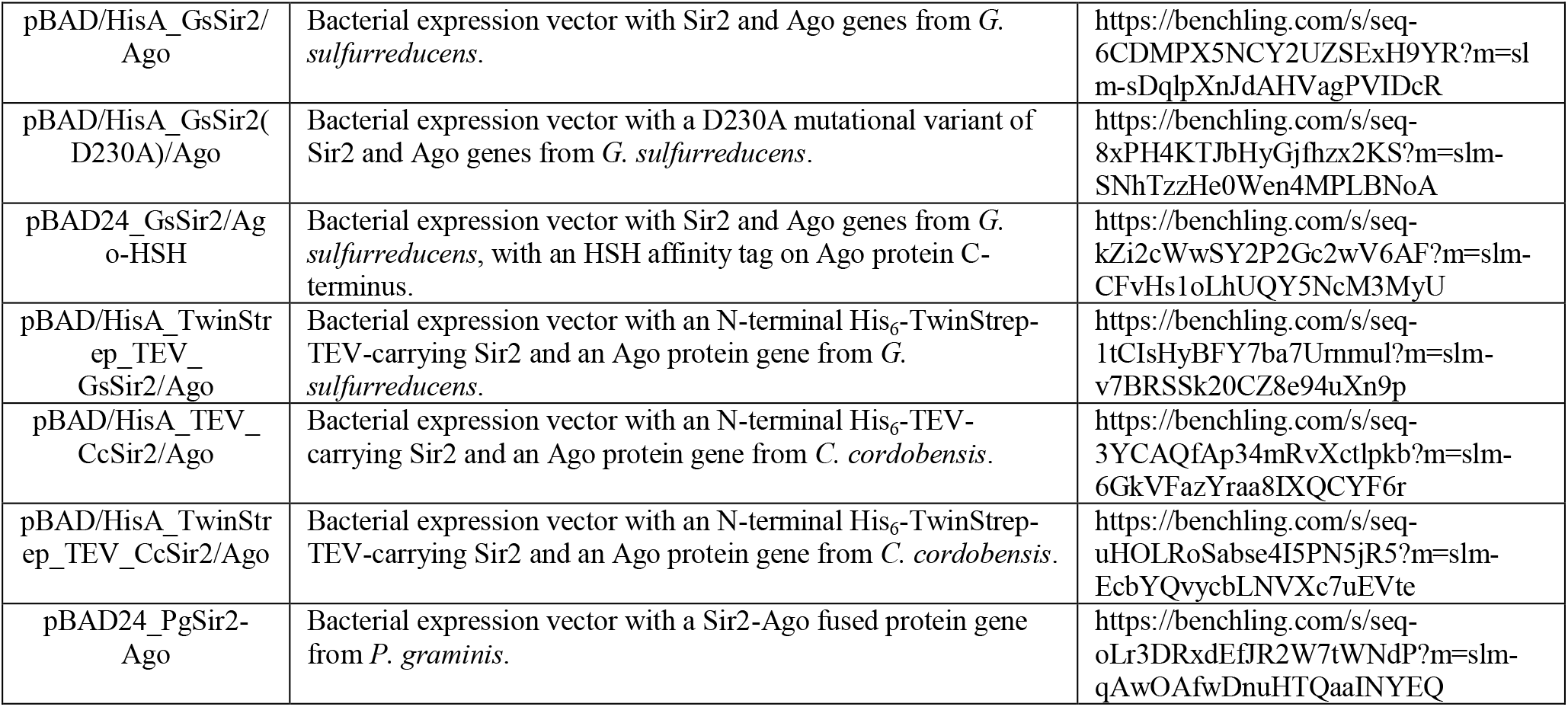
Strains, bacteriophages, plasmids, and proteins used in this work.

**Supplementary Table 2.**
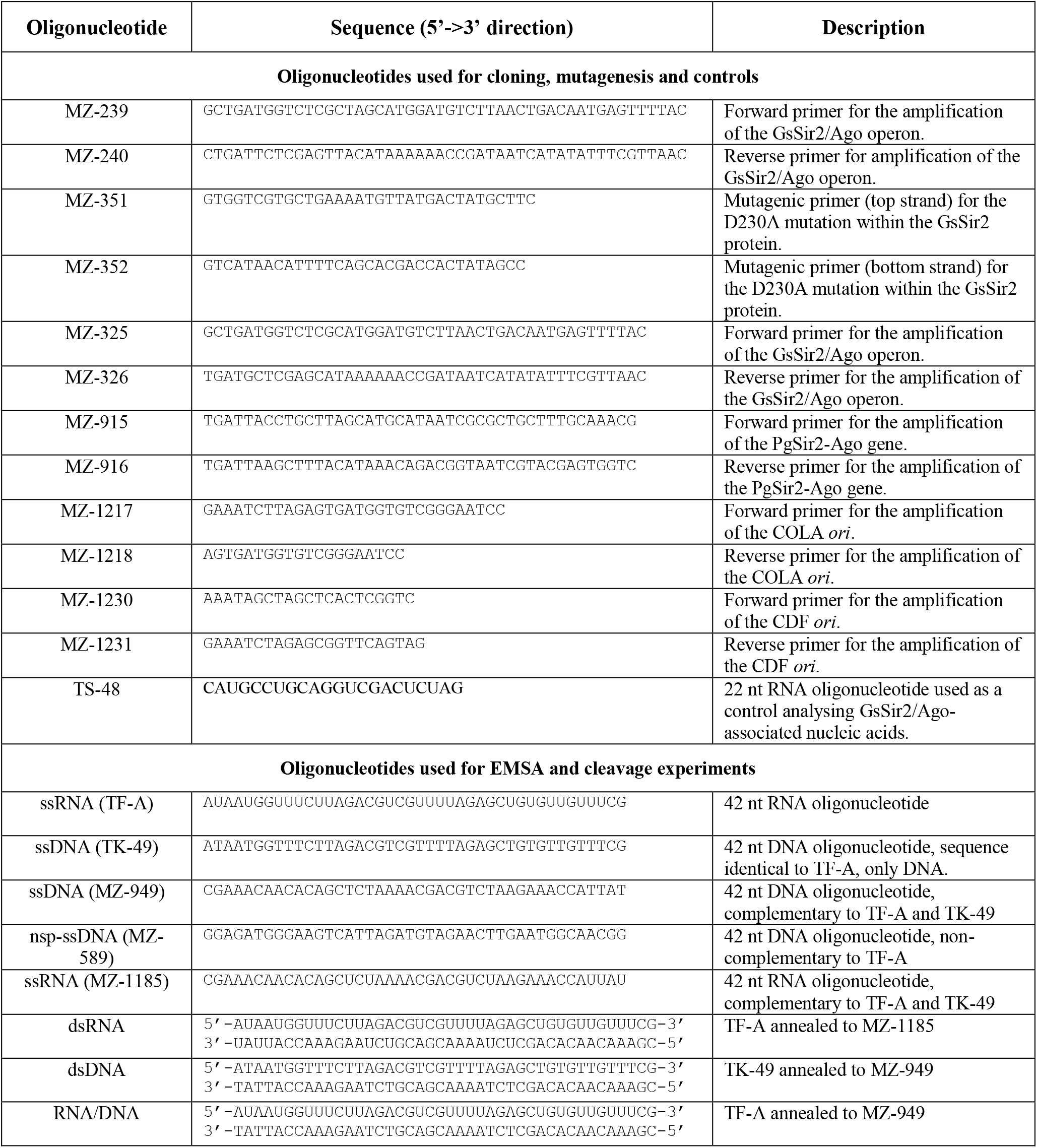
Oligonucleotides used in this work.

**Supplementary Table 3.** Parameters of the SAXS data.

| *Reciprocal space parameters* | *Merged SAXS normalized against water scattering merged data* |
| --- | --- |
| s min (nm<sup>-1</sup>) as in Guinier range | 0.1116200 |
| Guinier range (points) used for Guinier analysis | 48 to 170 |
| Rg, by Guinier analysis / AUTORG, nm | 3.61 / 3.61±0.034 |
| I(0) AUTORG, (cm<sup>-1</sup>) | 0.0734±0.00022 |
| Rg*s by Guinier analysis | 1.33281 |
| ***Molecular Mass estimations out of SAXS data, kDa*** |  |
| PRIMUS Qp | 107.9 |
| PRIMUS Vc | 99.7 |
| PRIMUS Size&Shape | 125.3 |
| PRIMUS Absolute scale | 101.1 |
| PRIMUS Bayesian inference | 101.1 |
| SAXSMoW server, v2.1 | 124.8 |
| ***Real-space parameters (GNOM)*** |  |
| s range (nm<sup>-1</sup>) used in GNOM | 0.1116 to 3.2449 |
| Dmax as parameter in GNOM / DATCLASS / SHANUM, nm | 11.4 / 12.3 / 13.4 |
| Rg real GNOM / DATPOROD, nm | 3.579±0.005044 / 3.58 |
| I(0) real GNOM / DATPOROD, (cm<sup>-1</sup>) | 0.07277±0.0001016 / 0.0728 |
| Porod volume DATPOROD (nm<sup>3</sup>) | 185.06 |
| SASDB ID | SASDNH2 |

